# Real-time single-molecule imaging in zebrafish embryos uncovers non-canonical translation

**DOI:** 10.64898/2026.01.05.697684

**Authors:** Maëlle Bellec, Jie Liang, Kenny Mattonet, Tatsuya Morisaki, Margaux Lay, Hans-Martin Maischein, Damien Avinens, Vincent Martinet, Delphine Muriaux, Timothy J Stasevich, Jérémy Dufourt, Didier Y R Stainier

## Abstract

Precise spatiotemporal control of protein synthesis is essential during embryogenesis, yet directly measuring translation kinetics *in vivo* remains challenging in vertebrates. In particular, it remains unclear how translation efficiency is determined for key developmental regulators, and which kinetic steps limit their production. Here, we used ALFA array-based nascent chain labeling combined with lattice light-sheet microscopy to visualize *bmp2b* translation in real time and at single-molecule resolution in early zebrafish embryos. When combined with MS2/MCP labeling to visualize all *bmp2b* mRNAs, we found that only some of them are being actively translated, suggesting that limited mRNA translation competence contributes to overall translation efficiency. We found that *bmp2b* translation operates below a maximal initiation regime, as replacement of its UTRs with viral UTRs increases ribosome loading. Furthermore, the *bmp2b*, but not *actb2*, 5′UTR supports cap-independent translation with ribosome loading comparable to cap-dependent initiation. Altogether, this approach provides a quantitative *in vivo* framework to dissect translation kinetics during early vertebrate development.

**Teaser:** Single-molecule imaging reveals *bmp2b* translation dynamics and cap-dependent and - independent initiation in zebrafish embryos.

## Introduction

The execution of complex developmental programs relies on the precise regulation of gene expression in both space and time. While transcriptional profiling has provided valuable insight into mRNA abundance, it is increasingly recognized that mRNA levels do not always reflect protein output, especially during rapid cell state transitions such as those observed during embryogenesis (*1–3*). These observations highlight the importance of studying gene expression as a dynamic and multi-layered process, where regulation extends beyond transcription to encompass mRNA localization, translation, and degradation.

Among these processes, translational control provides a rapid and flexible mechanism to modulate protein production without altering transcript abundance. By adjusting the efficiency and timing of protein synthesis, translation can fine-tune gene expression dynamics during developmental processes such as cell fate specification and tissue patterning, as well as during regeneration (*3–10*). This level of regulation is especially relevant for morphogens whose spatial concentration gradients must be tightly controlled to ensure robust pattern formation (*8*, *11*, *12*). In zebrafish, the bone morphogenetic protein gene *bmp2b* encodes a central morphogen that establishes the dorsoventral axis and controls gastrulation (*13–16*). *bmp2b* transcripts are distributed in a ventral-to-dorsal gradient (*17*), yet surprisingly little is known about how their translation is regulated. Notably, despite Bmp2b’s critical role in early development, Bmp2b protein levels are low or absent from proteomic datasets (*18*, *19*), and ribosome profiling indicates low translation efficiency (*20*). These observations raise an intriguing question: how are Bmp2b protein levels quantitatively controlled to generate a robust morphogen gradient during early development?

One possibility is that translation of *bmp2b* mRNAs is regulated at multiple levels, including the fraction of mRNAs engaged in translation, as well as ribosome initiation and elongation rates. However, the real-time dynamics of *bmp2b* translation *in vivo* remain largely unexplored, limiting our understanding of how morphogen production is quantitatively controlled during early vertebrate development.

To investigate how Bmp2b translation is regulated, it is essential to consider the regulatory elements that influence translation, especially the 5′ cap, the poly(A) tail and untranslated regions (UTRs) of mRNAs. The 5′ cap promotes 40S ribosome recruitment and canonical initiation (*21–23*), while 5′UTRs contain structures and motifs that modulate the initiation and efficiency of translation (*24–27*). A striking example of such regulation is provided by viruses as for example the 5′UTR of the *Orthoflavivirus* Zika virus (ZIKV) RNA, which contains highly structured elements and has been shown to exhibit internal ribosome entry site (IRES) activity in cell culture (*28*, *29*). This IRES activity enables cap-independent translation, allowing viral protein synthesis to proceed even when host cap-dependent translation is compromised (*28*, *29*). Moreover, ZIKV can infect zebrafish (*30*), offering a unique opportunity to study these non-canonical translational mechanisms in a live vertebrate model. Although IRES-mediated and, more broadly, cap-independent translation are well characterized *in vitro* and in viral systems, their role in vertebrate genes remains poorly understood. This lack of knowledge is largely due to the complexity of *in vivo* regulatory environments and the technical challenges of monitoring translation dynamics with high spatial and temporal resolution in developing organisms.

Efforts to dissect translation dynamics *in vivo* have long been constrained by the limited availability of tools that can monitor translation at high resolution in living organisms. Antibody-based nascent chain imaging approaches have emerged as powerful strategies to visualize translation at the single-cell level (*31–36*) and more recently, have been successfully adapted to *Drosophila* embryos (*37–41*). However, their application in vertebrate systems remains at an early stage. Technical challenges, including context-dependent variability in protein stability, heterogeneous expression, and limitations in imaging depth, continue to hinder their widespread implementation in complex tissues (*36*, *42*). Overcoming these challenges in zebrafish embryos would represent a significant advance, enabling the study of translation in a model that offers greater spatial and developmental complexity than cell culture or invertebrate systems.

Here, we deploy the ALFA_array system to achieve real-time imaging of *bmp2b* translation in live zebrafish embryos, making the first application of this technology in a vertebrate system. We first generated stable transgenic lines expressing a nanobody-based reporter for nascent peptide detection and validated the system using synthetic mRNA and plasmid injections. Using Lattice Light Sheet microscopy (*43*), we tracked individual *bmp2b* polysomes with subcellular resolution. We then generated a stable transgenic line expressing tagged *bmp2b* to quantify its native translation dynamics, and estimated its kinetics parameters. Furthermore, by exploiting the unique properties of ZIKV UTRs, we were able to probe both cap-dependent and cap-independent mechanisms in a physiological context.

Altogether, this work expands the translational imaging toolbox for vertebrate embryos and provides a framework to quantitatively study mRNA translation *in vivo*, including the respective contributions of translation competence, ribosome initiation rates, and UTR-dependent regulatory mechanisms.

## Results

### Implementation of the ALFA_array system for *in vivo* monitoring of translation

To establish the first live imaging approach for visualizing translation in zebrafish embryos using antibody-based detection of nascent peptides, we first adapted the well-established SunTag system (*31*, *32*, *34*, *35*), a gold standard for monitoring translation dynamics in live cells and *Drosophila* embryos through single-chain variable fragment (scFv) recognition of epitope-tagged nascent chains. We generated a transgenic zebrafish line expressing an anti-suntag scFv fused to monomeric superfolder GFP2 (scFv-msGFP2) (*44*), a fluorescent protein previously shown to display high brightness and monomericity in *Drosophila* (*38*), under the control of a ubiquitin promoter (*45*). Despite successful transgene expression, we observed prominent fluorescent aggregates beginning at 8 hours post-fertilization (hpf) (fig. S1A), limiting the utility of this system for live imaging of translation in zebrafish.

To overcome the limitations encountered with the SunTag system, we next implemented an orthogonal approach, originally developed in *Drosophila* embryos and mammalian cells, the ALFA_array system (*37*) (Fig. 1A). This bipartite strategy comprises multiple copies of the ALFA-tag (*46*) inserted into the gene of interest and specifically recognized with high affinity (*46*) by a fluorescently labeled anti-ALFA-tag nanobody (NB-ALFA). To enable broad applicability of this method in zebrafish, we generated a transgenic line that ubiquitously expresses an optimized NB-ALFA fused to msGFP2 (see methods) (Fig. 1A).

**Fig. 1.**
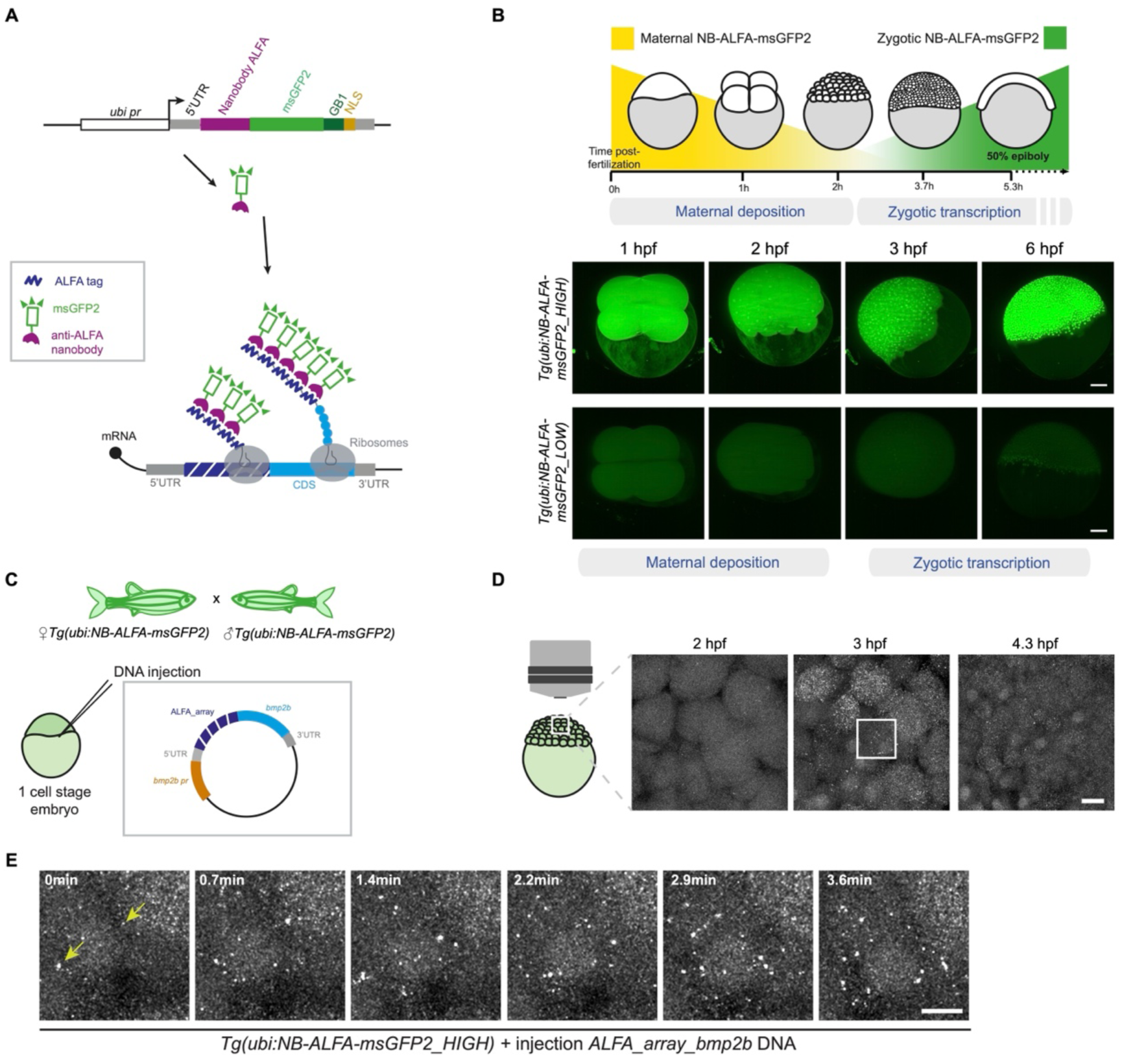
Implementation of the ALFA_array system in zebrafish embryos for live imaging of translation. (**A**) Schematic of the ALFA_array system for live imaging of translation. On top, schematic of the construct used to generate transgenic lines expressing NB-ALFA-msGFP2 under the control of a ubiquitin promoter. Upon translation, ALFA tags (dark blue) inserted into the gene of interest (blue, CDS: coding sequence) become bound by the anti-ALFA nanobody (NB-ALFA) fused to a fluorescent protein (green); GB1, Immunoglobulin-binding protein G, B1 domain; NLS, nuclear localization signal. (**B**) Schematic representation of the maternal-to-zygotic transition in zebrafish with the maternal deposition of NB-ALFA-msGFP2 in yellow and the zygotic NB-ALFA-msGFP2 expression in green. Below, maximum intensity projected images of light sheet movies of *Tg(ubi:NB-ALFA-msGFP2_HIGH)* and *Tg(ubi:NB-ALFA-msGFP2_LOW)* embryos with maternal and zygotic expression of the anti-ALFA nanobody. (**C**) Schematic of the cross used to live image translation in zebrafish embryos as shown in (D). Below, schematic of the plasmid containing *bmp2b* fused to the ALFA_array tag, which was injected into one-cell stage *Tg(ubi:NB-ALFA-msGFP2_HIGH)* embryos. (**D**) Schematic of the experimental set-up to live image translation in a zebrafish embryo (left) using confocal microscopy. Maximum intensity projected images from a confocal movie of a *Tg(ubi:NB-ALFA-msGFP2_HIGH)* embryo injected with *ALFA_array_bmp2b* DNA (right). White square represents the zoomed area for the images shown in (E). (**E**) Zoomed in images from the confocal movie in (D). Yellow arrows point to translation dots. Scale bars: 100 μm (B), 20 μm (D), 10 μm (E).

We first observed different expression levels of the NB-ALFA-msGFP2 transgene in the F1 generation (Fig. S1B-C) and established two lines, one with “low expression” (*Tg(ubi:NB-ALFA-msGFP2_LOW)*) and one with “high expression” (*Tg(ubi:NB-ALFA-msGFP2_HIGH)*), the latter showing a ∼2.5 fold increase in expression compared with the former (fig. S1D). Next, by using light sheet microscopy, we confirmed that NB-ALFA-msGFP2 was maternally deposited (Fig. 1B, Movie S1) and started to be zygotically expressed at around 4 hpf (fig. S1E-F, Movie S1). Importantly, light sheet and confocal imaging of these reporter lines revealed diffuse and uniform fluorescence during early development, with no evidence of aggregation (Fig. 1B and S1G), thereby enabling their use for high-resolution live imaging of translation.

To investigate the translation dynamics of a key developmental gene, we applied the ALFA_array system to the morphogen Bmp2b. We inserted an array of 32x ALFA-tags at the N-terminus of Bmp2b and injected *in vitro* transcribed mRNA into one-cell stage wild-type (WT) embryos (fig. S2A). Western blot analysis detected a predominant band at ∼115 kDa, corresponding to the molecular weight expected for the full-length fusion protein (ALFA_array_Bmp2b) (fig. S2B); it also revealed ALFA_array_Bmp2b expression as early as 1.5 hours post-fertilization (hpf), followed by a substantial decrease by 6 hpf (fig. S2B). No signal was detected in embryos injected with *bmp2b* mRNA lacking the ALFA_array, thereby confirming the specificity and low background of the detection system (fig. S2B). To better mimic endogenous regulation, we next injected a plasmid containing the 2.27 kb *bmp2b* promoter (*16*) and intronic sequences (fig. S2C), which drove expression of ALFA-tagged Bmp2b from approximately 4.3 hpf, with levels increasing at least until 12 hpf (fig. S2D). This expression reflects the temporal pattern of endogenous *bmp2b* mRNA (*47*) (fig. S2E) and is consistent with ribosome profiling data (*48*) (fig. S2F).

To determine whether the *bmp2b-ALFA-array* construct retains biological activity and whether NB-ALFA binding interferes with its function, we performed functional assays *in vivo*. We injected the *bmp2b-ALFA-array* construct into both wild-type embryos and embryos expressing NB-ALFA-msGFP2, and compared the resulting phenotypes with those obtained after injection of non-tagged *bmp2b* mRNA, which is well known to induce ventralization (*16*, *49*, *50*). In both genetic backgrounds, injection of the *bmp2b-ALFA-array* construct into one-cell stage embryos produced ventralization phenotypes comparable to those observed in embryos injected with the non-tagged *bmp2b* control (fig. S3A), indicating that the ALFA_array does not abolish Bmp2b function and that NB-ALFA binding does not measurably affect its biological activity.

To further test the binding properties of NB-ALFA in zebrafish embryos, we performed Fluorescence Recovery After Photobleaching (FRAP) in *Tg(ubi:NB-ALFA-msGFP2_HIGH)* embryos injected with an *H2B_mCherry_ALFA-tag* plasmid (*51*) (fig. S2G). We observed minimal recovery of NB-ALFA-msGFP2 fluorescence over a 4.5-minute period (fig. S2, G and H), consistent with strong binding of the nanobody to the ALFA-tag. This finding aligns with previously published data showing slow recovery kinetics of H2B, which typically occurs over hours (*51*, *52*). In contrast, FRAP analysis in embryos expressing only NB-ALFA-msGFP2 (i.e., without an ALFA-tagged target) revealed rapid recovery, consistent with the free diffusion of the unbound nanobody as well as the absence of visible aggregates (fig. S2I).

Next, to visualize translation in zebrafish embryos, we initially performed conventional live confocal imaging on *Tg(ubi:NB-ALFA-msGFP2_HIGH)* embryos injected with the ALFA_array_*bmp2b* plasmid (Fig. 1C). From approximately 2.5 hpf, we observed the emergence of discrete fluorescent foci in a mosaic pattern (Fig. 1D), consistent with specific translation events. We observed that these puncta were motile, likely reflecting active sites of *bmp2b* translation (Fig. 1E).

In summary, we have established transgenic zebrafish lines that ubiquitously express NB-ALFA throughout development, providing a robust, aggregation-free platform for antibody-based detection of nascent peptides *in vivo*. Using the *ALFA_array_bmp2b* reporter, we directly visualize active *bmp2b* translation in early zebrafish embryos by detecting the stable binding of NB-ALFA-msGFP2 to ALFA-tagged nascent chains, an essential requirement for accurate and quantitative analysis of mRNA translation dynamics.

### Real-time visualization of *bmp2b* translation dynamics on single mRNAs in zebrafish embryos using Lattice Light Sheet microscopy

Although translation sites were clearly detectable with confocal imaging, the temporal resolution was insufficient for detailed kinetic analysis, and the translation spots could not be resolved at higher frame rates. To overcome these limitations, we employed Lattice Light Sheet Microscopy (LLSM) for high-resolution, rapid volumetric imaging with minimal phototoxicity (Fig. 2A). Using this imaging technique, we achieved volumetric imaging over an area of approximately 300 × 100 microns with a depth of 35 microns at a rate of one stack per second, enabling real-time tracking of translation foci over several minutes across multiple cells simultaneously (Fig. 2, B and C, Movie S2). We detected bright foci, which were absent in cells without *ALFA_array_bmp2b* expression, likely corresponding to translation sites (polysomes) (fig. S3B). To test whether these foci indeed represented active *bmp2b* translation, we performed control injections with either *ALFA_array_bmp2b* mRNA, *ALFA_array_bmp2b* mRNA together with the translation inhibitor puromycin, or *bmp2b* mRNA lacking the ALFA_array. Fluorescent foci were completely absent in both puromycin-treated embryos and ALFA-less conditions (Fig. 2D, Movie S3), implying that the observed signal arises from active translation sites.

**Fig. 2.**
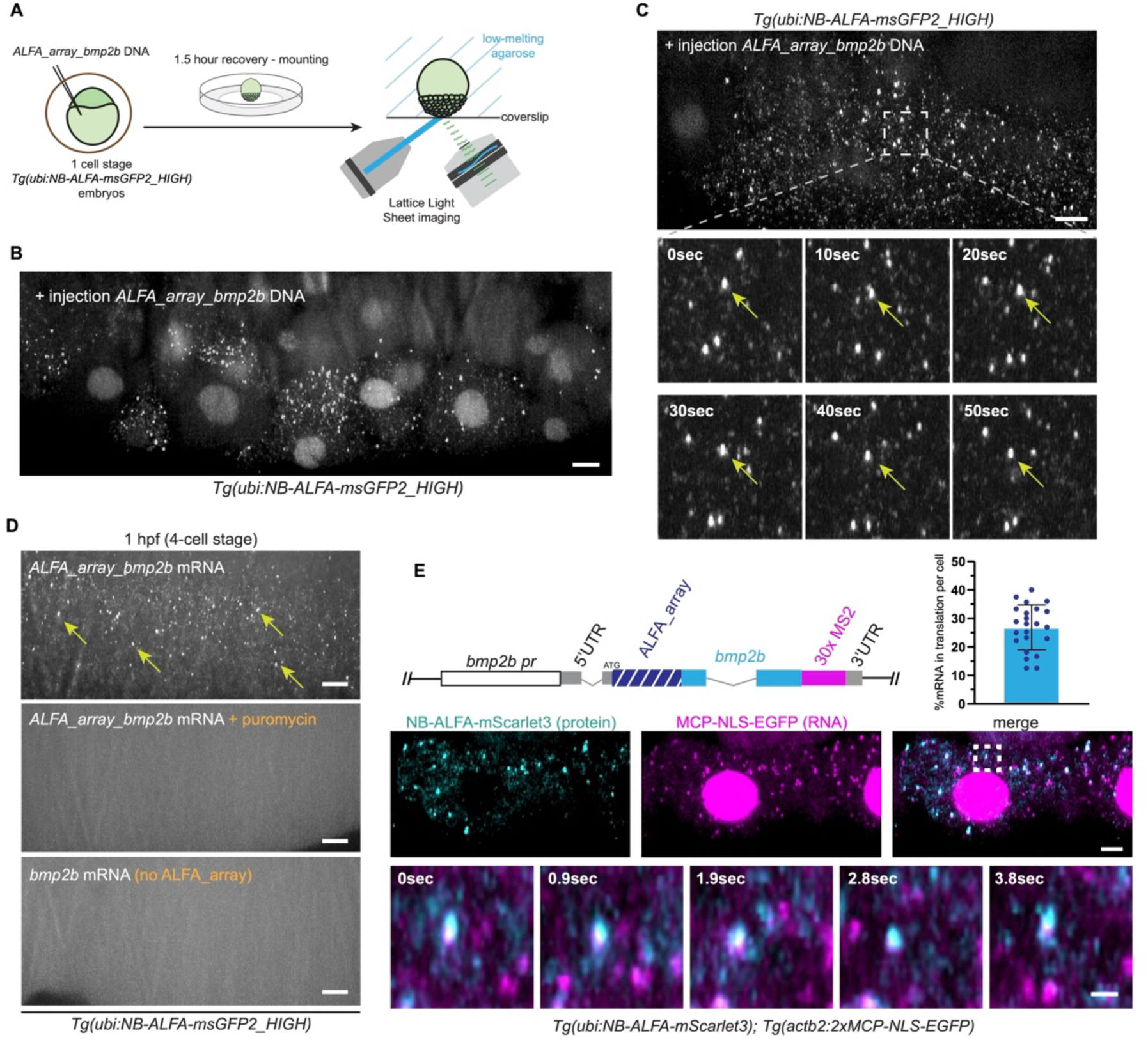
Visualization and validation of *bmp2b* translation in zebrafish embryos using Lattice Light-Sheet microscopy. (**A**) Schematic of experimental set-up for LLS imaging of zebrafish embryos. Embryos were injected at the one-cell stage, manually dechorionated and then mounted in low melting point agarose in a glass-bottomed dish with the animal pole oriented toward the coverslip. (**B**) Maximum intensity projected image from an LLS movie of a ∼4.3 hpf *Tg(ubi:NB-ALFA-msGFP2_HIGH)* embryo injected with *ALFA_array_bmp2b* DNA. (**C**) Maximum intensity projected image from an LLS movie of a ∼4.3 hpf *Tg(ubi:NB-ALFA-msGFP2_HIGH)* embryo injected with *ALFA_array_bmp2b* DNA, and zoomed in images from the movie showing the translation dots of *ALFA_array_bmp2b* (arrow) over time. (**D**) Representative maximum intensity projected images from LLS movies of 1 hpf *Tg(ubi:NB-ALFA-msGFP2_HIGH)* embryos injected with *ALFA_array_bmp2b* mRNA without or with puromycin or with *bmp2b* mRNA (no ALFA_array). Yellow arrows point to translation dots, absent in the presence of puromycin or when embryos were injected with the construct lacking the *ALFA_array*. (**E**) Schematic of the *bmp2b* construct containing the *ALFA_array* and 30x MS2. Representative image from an LLS movie of a ∼4.3 hpf *Tg(ubi:NB-ALFA-mScarlet3); Tg(actb2:2xMCP-NLS-EGFP)* embryo injected with *ALFA_array_bmp2b_30xMS2* DNA. MCP-EGFP is represented in magenta and corresponds to mRNA molecules (nuclear accumulation is due to the presence of NLS), and NB-ALFA-mScarlet3 (nascent translation) is represented in cyan. Histogram shows the quantification of the percentage of mRNA molecules in translation (colocalized MCP-NLS-EGFP and NB-ALFA-mScarlet3 signals). Scale bars: 10 μm (B, C, D, E), 5 μm (E), 1 μm (zoomed in panel E).

We first validated the ALFA_array system in fixed embryos, using a mosaic expression approach through plasmid injection, with Hybridization Chain Reaction Fluorescence In Situ Hybridization (HCR-FISH) for ALFA-tagged *bmp2b* mRNA and immunostaining for the ALFA tag epitopes. We observed clear co-expression (fig. S3C), thereby demonstrating specificity and active translation of the mRNAs generated from the injected plasmid. However, this approach in fixed embryos fails to achieve reliable single-molecule resolution, thereby preventing quantitative measurement of translation of individual mRNAs or the kinetic analysis of Bmp2b protein output.

To overcome this limitation, we next aimed to simultaneously visualize single mRNAs and their translation in live. To this end, we combined the ALFA_array system with the MS2/MCP system (*53*) by generating a construct containing *ALFA_array_bmp2b* and 30 MS2 loops in the 3’UTR of *bmp2b* (Fig. 2E). We then established a zebrafish line stably expressing the nanobody fused to mScarlet3, crossed it with the *Tg(actb2:2xMCP-NLS-EGFP)* line (*54*), and injected into the resulting progeny the tagged *bmp2b* construct. This strategy, together with the high temporal resolution of lattice light-sheet microscopy (180 ms/frame), allowed us to visualize putative transcription sites in nuclei, as identified by their larger size, higher fluorescence intensity, and more restricted movement compared with individual mRNA molecules (*37*, *54*) or individual mRNA molecules in motion (fig. S3D). Importantly, we could also monitor translation occurring on these single mRNA molecules (Fig. 2E and Movie S4-5) and found that approximately 26% of mRNAs were actively translating (Fig. 2E). Consistent with reports that ∼20-60% of mRNAs from various genes are being actively translated in live vertebrate cells, our data place *bmp2b* within the modestly translated range characteristic of regulated transcripts (*32*, *34*).

### Stable insertion of an ALFA_array-tagged *bmp2b* transgene enables *in vivo* quantification of translation kinetics in developing zebrafish embryos

To analyze translation dynamics in a more physiologically relevant context, we generated a stable transgenic zebrafish line expressing *bmp2b* tagged with the ALFA_array and under the control of the 2,27 kb *bmp2b* promoter (*Tg(bmp2b:ALFA_array_bmp2b)*) (Fig. 3A). This line recapitulated the expression pattern reported for the proximal 2,27 kb *bmp2b* promoter (*16*, *55*), exhibiting broad expression at 4.3 hpf and restricted expression in the dorsal organizer and to a lesser extent on the ventral side by shield stage (6 hpf) (Fig. 3B). We further confirmed the specificity of the translation foci by their absence outside the *bmp2b* transgene expression domain (fig. S4A). Time-lapse imaging of *bmp2b* translation from 4.3 to 6.5 hpf revealed a pattern initially broad across the embryo and later restricted to one side (fig. S4B). We also observed *bmp2b* mRNA translation in the developing eye at 24 hpf using LLS imaging (fig. S4C).

**Fig. 3.**
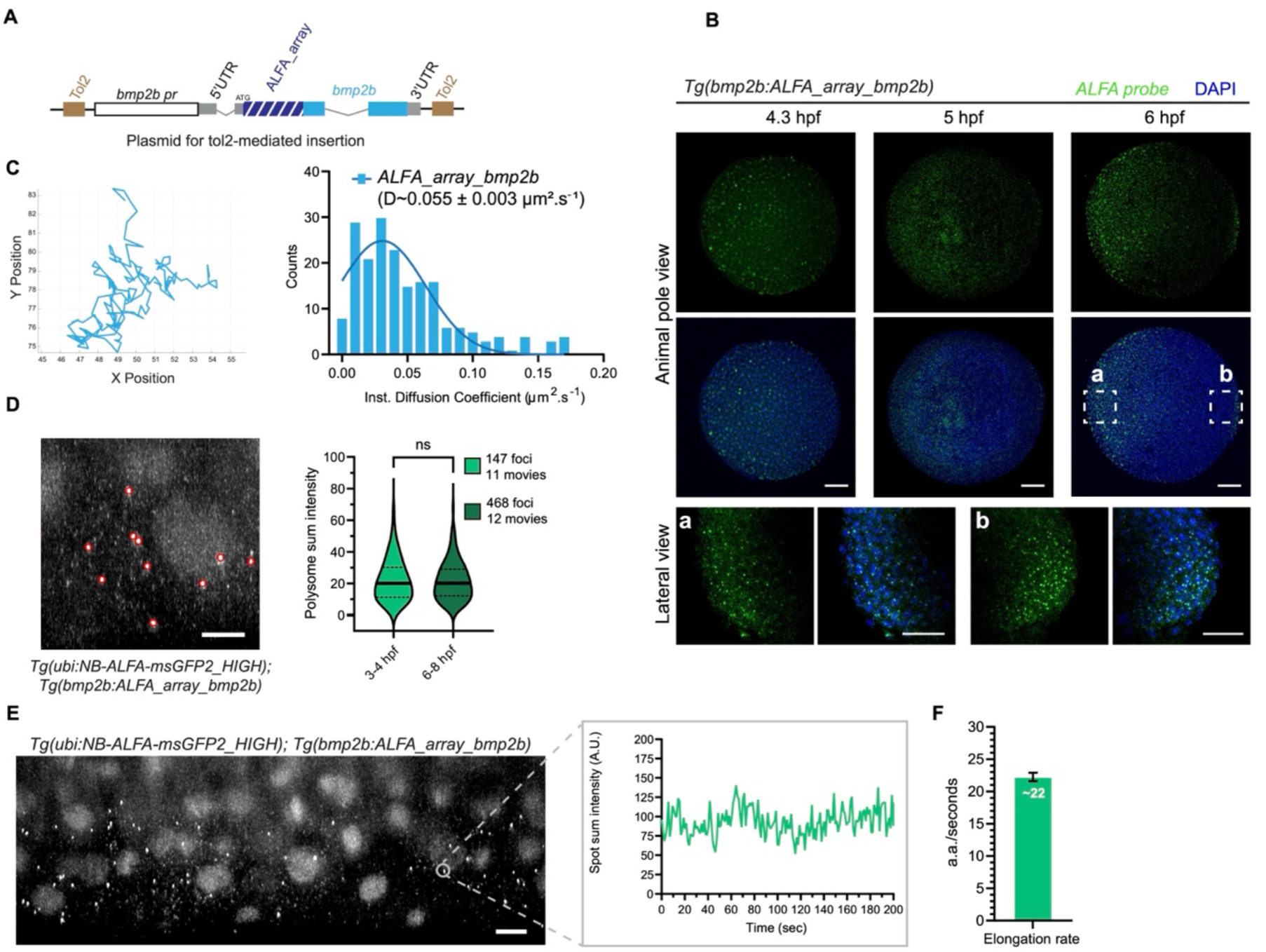
Tracking and quantifying *bmp2b* polysomes to determine translation kinetics. (**A**) Schematic of the *bmp2b* transgene construct containing the *ALFA_array* used for tol2-mediated insertion. (**B**) Images from fluorescence in situ hybridization using a probe against *ALFA_array* mRNA (green) in *Tg(bmp2b:ALFA_array_bmp2b)* embryos in animal pole view; ventral to the left. Bottom, zoomed in images show the transgene expression at 6 hpf in the ventral (a) and dorsal (b) side. (**C**) On the left, representative trajectory of a *bmp2b* mRNA particle in translation; x and y positions are in μm. On the right, frequency distribution of instantaneous diffusion coefficient of *bmp2b* mRNA molecules in translation in 3-4.3 hpf *Tg(ubi:NB-ALFA-msGFP2_HIGH); Tg(bmp2b:ALFA_array_bmp2b)* embryos (n=259 traces, from 12 movies). D is the mean diffusion coefficient with the standard error of the mean (SEM). (**D**) On the left, representative maximum intensity projected image of polysome detection (red circles) from an LLS movie of a ∼6 hpf *Tg(ubi:NB-ALFA-msGFP2_HIGH); Tg(bmp2b:ALFA_array_bmp2b)* embryo. On the right, polysome sum intensity quantification (see methods) from 3-4.3 hpf (n=147 from 11 movies) and 6-8 hpf (n=468 from 12 movies) embryos. Black line represents the median; ns: non-significant with unpaired *t* test (p-value >0.05). (**E**) On the left, representative maximum intensity projected image from an LLS movie of a ∼6 hpf *Tg(ubi:NB-ALFA-msGFP2_HIGH); Tg(bmp2b:ALFA_array_bmp2b)* embryo and, on the right, example of the intensity over time of a tracked *bmp2b* polysome (grey circle and inset). (**F**) Mean values of the elongation rate of *ALFA_array_bmp2b* mRNA during translation (n=43). The estimated parameter is indicated with the respective 95% confidence intervals. Scale bars: 100 μm (B), 50 μm (zoomed in panels B), 5 μm (D), 10 μm (E).

We then analyzed published ribosome profiling and RNA-seq datasets (*27*, *56*) and found that *bmp2b* belongs to a group of genes with low translation efficiency (fig. S4, D and E). To better understand the basis for this low efficiency, we sought to determine the dynamic features of *bmp2b* translation directly and *in vivo*. We first tracked individual translating *bmp2b* mRNAs (Fig. 3C), calculated their instantaneous diffusion coefficient, and compared *ALFA_array_bmp2b* expression originating from the injected plasmid with that from the stably integrated transgene. Polysome trajectories in both experimental conditions showed no preferential directional movement (fig. S5A), and mean square displacements as well as diffusion coefficients were comparable between mRNAs originating from the *ALFA_array*_*bmp2b* plasmid (fig. S5, B and C) and the transgene (Fig. 3C and fig. S5D), with an average diffusion rate of ∼0.055 μm²/s (Fig. 3C and fig. S5C), comparable to measurements reported for mRNA diffusion in invertebrate and mammalian systems (*32*, *37*). Additionally, we quantified polysome intensities at single time points, which provides a readout of the number of nascent polypeptide chains on each mRNA. Since each actively translating ribosome contributes to the fluorescence signal of the ALFA_array, these intensities serve as a proxy for ribosome occupancy on the mRNA. Using this approach, we observed that at the single-molecule level *bmp2b* mRNAs were translated with similar efficiency both before and during gastrulation (3–4.3 hpf and 6–8 hpf, respectively; Fig. 3D), indicating stable ribosome loading across these developmental stages. Background fluorescence levels in translating cells remained unchanged (fig. S5E), indicating that the pool of free NB-ALFA remains stable between the analyzed time points in *Tg(bmp2b:ALFA_array_bmp2b)* embryos. Furthermore, we observed an increased number of foci per cell at 6-8 hpf compared with 3-4.3 hpf, reflecting the higher mRNA expression level at this stage, as measured by RNA-sequencing (fig. S5F).

We next sought to determine the kinetics of translation on single mRNAs. Using LLSM to track individual polysome foci in 3-6 hpf embryos and extracted single-spot intensity fluctuations as a quantitative readout of translation dynamics (*33*, *37*, *57*) (Fig. 3E, Movie S6). After applying photobleaching correction (see Methods and fig. S5G), and autocorrelation analysis and fitting (fig. S5, H to J), we estimated an average elongation rate of ∼22 amino acids (a.a.) per second Fig. 3F. While this elongation rate estimate may represent an upper bound and does not represent the heterogeneity between transcripts, it is within the range of previously reported rates in mammalian cells (from ∼3 to 22 a.a. per second) (*31–33*, *58–61*) and *Drosophila* embryos (∼5 and 35 a.a. per second) (*37*, *41*).

Altogether, these findings establish the ALFA_array system as a physiologically relevant platform for real-time studies of translation dynamics in vertebrate embryos.

### The poly(A) tail promotes *bmp2b* translation with moderate impact on ribosome loading

In zebrafish embryos, the poly(A) tail is essential for post-transcriptional control, allowing maternal mRNAs to be properly stored, activated, and degraded in a tightly regulated developmental program (*62*, *63*). Although poly(A) tails are well known to enhance translation by promoting ribosome recruitment (*64*, *65*) and increasing mRNA stability (*63*), it remains difficult to distinguish between these two contributions in the context of a living embryo. Moreover, the relationship between poly(A) tail length and translation efficiency is context-dependent and does not always show a direct correlation (*62*).

To investigate how the presence of a poly(A) tail influences *bmp2b* mRNA translation in the early zebrafish embryo, we compared by western blot analysis the protein output following *bmp2b* (tagged with the ALFA_array) mRNA injection with and without a poly(A) tail. We observed that non-polyadenylated *bmp2b* mRNAs yielded lower protein output than their polyadenylated counterparts (fig. S6A). To determine whether this reduction in protein output was due to a decreased amount of mRNA competent for translation or impaired translation efficiency, we generated a chimeric construct by replacing the *bmp2b* 3′UTR with the ZIKV 3’UTR (Fig. 4A). Indeed, the ZIKV 3′UTR lacks a poly(A) tail but its RNA escapes degradation due to its highly structured conformation (*66*, *67*), thereby allowing one to uncouple mRNA stability from poly(A)-dependent translation. We *in vitro* transcribed mRNAs with either the *bmp2b* 3’UTR or the ZIKV 3’UTR, both without a poly(A) tail, and injected them in *Tg(ubi:NB-ALFA-msGFP2_HIGH)* embryos. LLS live imaging revealed that a higher proportion of mRNAs containing the ZIKV 3′UTR were engaged in translation (Fig. 4C) and that these mRNAs displayed higher polysome intensity, thereby resulting in increased protein output (fig. S6B). In contrast, *bmp2b* 3’UTR transcripts lacking a poly(A) tail displayed only a moderate decrease in polysome intensity compared with *bmp2b* 3’UTR with a poly(A) tail (Fig. 4B). Together, these results indicate that, for the *bmp2b* reporter, the poly(A) tail has only a modest effect on ribosome loading, suggesting that it primarily acts on the mRNA’s competence to be translated.

**Fig. 4.**
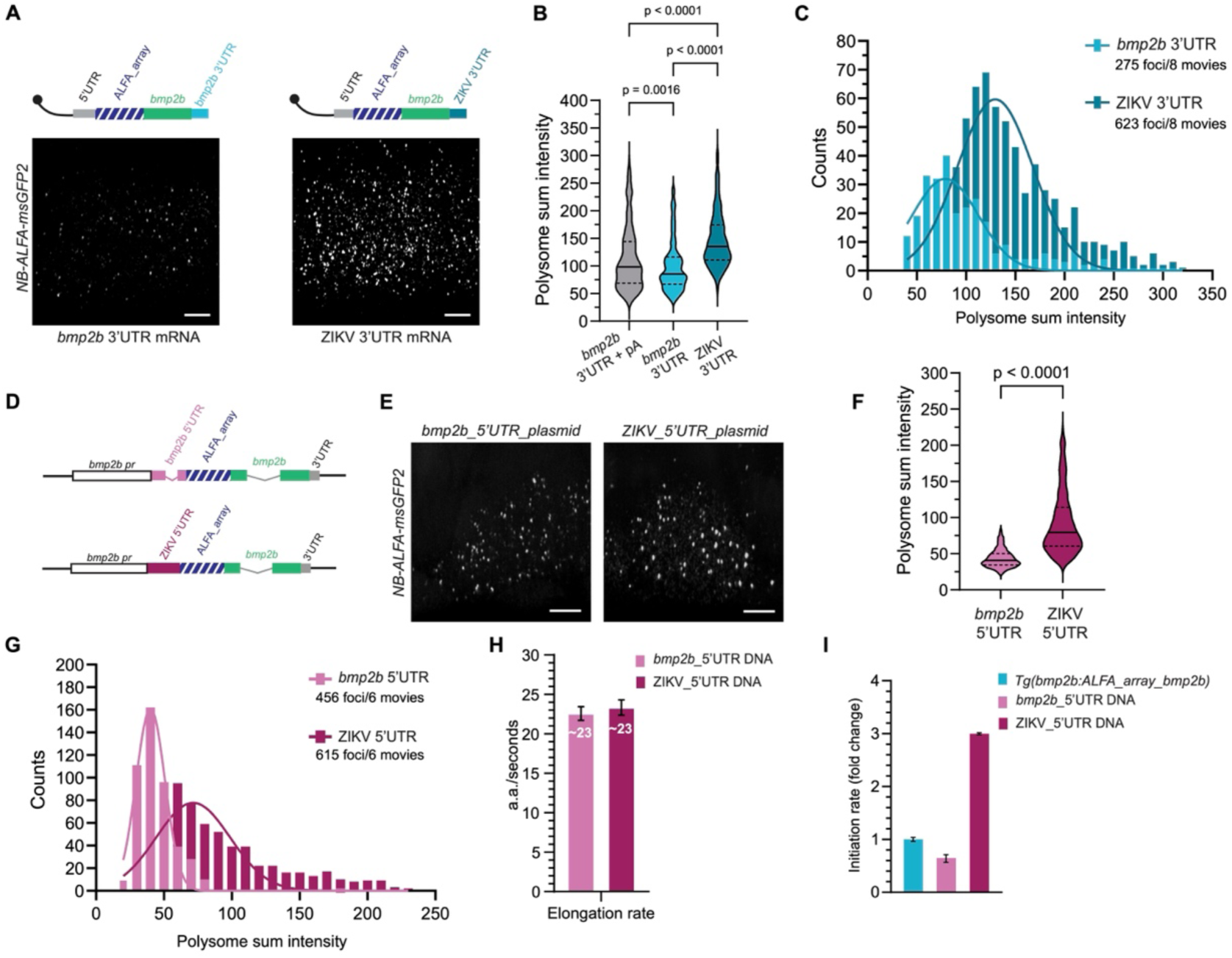
Comparative analysis of *bmp2b* and viral UTRs reveals *bmp2b* translation regulation in early zebrafish embryos. (**A**) Schematic of the *in vitro* transcribed mRNA of *ALFA_array_bmp2b* containing the *bmp2b* 3’UTR (light blue) or the ZIKV 3’UTR (dark blue) without poly(A) tail. Representative maximum intensity projected images from LLS movies of ∼1.5 hpf *Tg(ubi:NB-ALFA-msGFP2_HIGH)* embryos injected with the respective mRNA. (**B-C**) Violin plot (B) and frequency distribution (C) of polysome sum intensity of *bmp2b* 3’UTR (n= 275 from 8 movies) and ZIKV 3’UTR (n= 623 from 8 movies) *ALFA_array_bmp2b* mRNAs without poly(A) tail measured from the movies shown in (A). One-way ANOVA test was performed. Gaussian fitting is shown with the curves. (**D**) Schematic of the *ALFA_array_bmp2b* DNA containing the *bmp2b* 5’UTR (pink) or the ZIKV 5’UTR (dark pink). (**E**) Representative maximum intensity projected images from LLS movies of ∼4.3 hpf *Tg(ubi:NB-ALFA-msGFP2_HIGH)* embryos injected with the *bmp2b* 5’UTR or ZIKV 5’UTR *ALFA_array_bmp2b* DNA (represented in D). (**F**) Polysome sum intensity of *bmp2b* 5’UTR (n= 456 from 6 movies) and ZIKV 5’UTR (n= 615 from 6 movies) *ALFA_array_bmp2b* mRNAs measured from the movies shown in (C). Two-tailed Welch’s *t* test was performed. (**G**) Frequency distribution of individual polysome sum intensity from ∼1.5 hpf *Tg(ubi:NB-ALFA-msGFP2_HIGH)* embryos injected with the *bmp2b* 5’UTR or ZIKV 5’UTR *ALFA_array_bmp2b* DNA. Gaussian fitting is shown with the curves. (**H**) Mean values of the elongation rate of *bmp2b* 5’UTR (n= 45 from 6 movies) and ZIKV 5’UTR (n= 51 from 6 movies) *ALFA_array_bmp2b* mRNAs during translation measured from ∼4.3 hpf *Tg(ubi:NB-ALFA-msGFP2_HIGH)* embryos injected with the respective mRNAs. The estimated parameters are indicated with their respective 95% confidence intervals. (**I**) Mean ribosome initiation rates of *ALFA_array_bmp2b* mRNAs from *Tg(bmp2b:ALFA_array_bmp2b)* embryos (n = 43 from 11 movies), and from embryos injected with *bmp2b* 5′UTR (n = 45 from 6 movies) or ZIKV 5′UTR (n = 51 from 6 movies) *ALFA_array_bmp2b* DNA constructs, measured at ∼4.3 hpf in *Tg(ubi:NB-ALFA-msGFP2_HIGH)* embryos. Values are represented as fold change compared with the initiation rate of *ALFA_array_bmp2b* mRNAs from *Tg(bmp2b:ALFA_array_bmp2b)* embryos set to one. The estimated parameters are indicated with their respective 95% confidence intervals. Scale bars: 10 μm.

### Translation from the *bmp2b* 5′UTR is under initiation-level control

To directly assess the ribosome loading potential of the *bmp2b* mRNA, we substituted its 5’UTR with the ZIKV 5′UTR in our *ALFA_array_bmp2b* reporter, while keeping the coding sequence unchanged, enabling a direct comparison of their translation efficiency (Fig. 4D). We performed LLS imaging of *Tg(ubi:NB-ALFA-msGFP2_HIGH)* embryos following the injection of ALFA_array-tagged *bmp2b* plasmids containing either the *bmp2b* 5′UTR or the ZIKV 5′UTR under the same promoter, leading to the expression of endogenously capped and poly(A) tailed mRNAs (Fig. 4, D and E). We measured polysome intensities and observed that the ZIKV 5’UTR polysomes exhibit a higher intensity compared with the *bmp2b* 5’UTR polysomes (Fig. 4, F and G). Consistent with our single-molecule data, we observed increased protein output from the construct containing the ZIKV 5’UTR compared with the one containing the *bmp2b* 5’UTR (fig. S6C). We also observed increased protein output from *in vitro*-transcribed mRNAs when using the ZIKV 5’UTR (fig. S6D) compared with the *bmp2b* 5’UTR, indicating that the increase observed when injecting the plasmids is not due to differences in plasmid transcription levels. Altogether, these data suggest that ZIKV 5′UTR mRNAs may be covered by more ribosomes than *bmp2b* 5′UTR mRNAs, potentially reflecting differences in ribosome initiation rates. To test this hypothesis, we tracked individual polysomes and quantified temporal fluctuations in fluorescence intensity for both *bmp2b* and ZIKV 5′UTR constructs. Autocorrelation analysis of these signals (fig. S6, E and F) allowed extraction of translation kinetic parameters (Fig. 4H). As expected, elongation rates were similar between constructs (Fig. 4H; also consistent with the transgene in Fig. 3F), whereas ribosome initiation rates were up to fivefold higher on ZIKV 5′UTR transcripts (Fig. 4I). These results indicate that, in the early embryonic context, the *bmp2b* 5′UTR does not drive translation at maximal capacity.

### The *bmp2b* 5′UTR enables non-canonical translation initiation in early zebrafish embryos

Given that it was described in cells in culture that the ZIKV 5’ and 3’UTRs contain highly structured RNA elements capable of promoting cap-independent translation (*29*, *68*), we hypothesized that the observed increase in translation efficiency was due to an IRES-like activity in its 5’UTR, independent of the rest of its genome. To test this hypothesis, we first *in vitro* transcribed separately the *bmp2b* and ZIKV 5′UTR, fused to *bmp2b* coding sequence tagged with the ALFA_array (fig. S7A), with the non-functional cap analog A(ppp)G and a poly(A) tail and injected the resulting mRNAs into embryos for LLS imaging (Fig. 5A). Remarkably, in the absence of a functional cap, the ZIKV 5′UTR still drove robust polysome formation and translation (Fig. 5A), consistent with efficient cap-independent initiation. Furthermore, transcripts containing the ZIKV 5’UTR produced more protein (fig. S7B) and exhibited higher polysome intensity compared with those carrying the *bmp2b* 5′UTR (Fig. 5B) in the presence of cap analogs. Despite analyzing an equivalent number of imaging datasets, transcripts with the ZIKV 5′UTR also yielded a greater number of active polysomes (Fig. 5C), suggesting enhanced ribosome recruitment as well as increased mRNA engaged in translation. Interestingly, injection of *in vitro* transcribed mRNAs containing the *bmp2b* 5’UTR with the non-functional cap analog A(ppp)G led to a measurable level of translation (Fig. 5A and fig. S7C). This clearly detectable protein output from A(ppp)G-capped mRNAs further suggests that early zebrafish embryos can support non-canonical translation initiation. To assess specificity, we also injected *bmp2b* mRNAs bearing the *actin beta 2* (*actb2*) 5′UTR and capped with the same non-functional A(ppp)G. In this case, we did not detect substantial protein production (Fig. 5D), although with a functional cap, the *actb2* 5’UTR produced at least as much protein as the *bmp2b* 5’UTR (Fig. S7D), suggesting that cap-independent translation is not a general feature of all 5’UTRs. Furthermore, to investigate the mode of cap-independent translation with *bmp2b* 5’UTR in the early zebrafish embryo, we performed LLS imaging of *bmp2b* reporter mRNAs carrying either the A(ppp)G- or G(ppp)G-cap (Fig. 5E). Due to extensive cell movements during early zebrafish embryogenesis (0.5-2 hpf), long-term tracking of individual mRNAs to extract translation kinetics was not feasible. Although polysome intensities were slightly higher for mRNA with the non-functional cap, the two transcripts were nearly identical (Fig. 5F), indicating no major difference in ribosome initiation rates for cap-dependent and cap-independent translation. Because polysome intensities were comparable between injected *ALFA_array_bmp2b* mRNAs and plasmid-derived transcripts (fig. S7E), the observed effects are unlikely to result from overexpression. Together, these findings indicate that early zebrafish embryos can sustain efficient translation with comparable initiation rates through both cap-dependent and cap-independent mechanisms, thereby revealing a gene-specific layer of translational regulation.

**Fig. 5.**
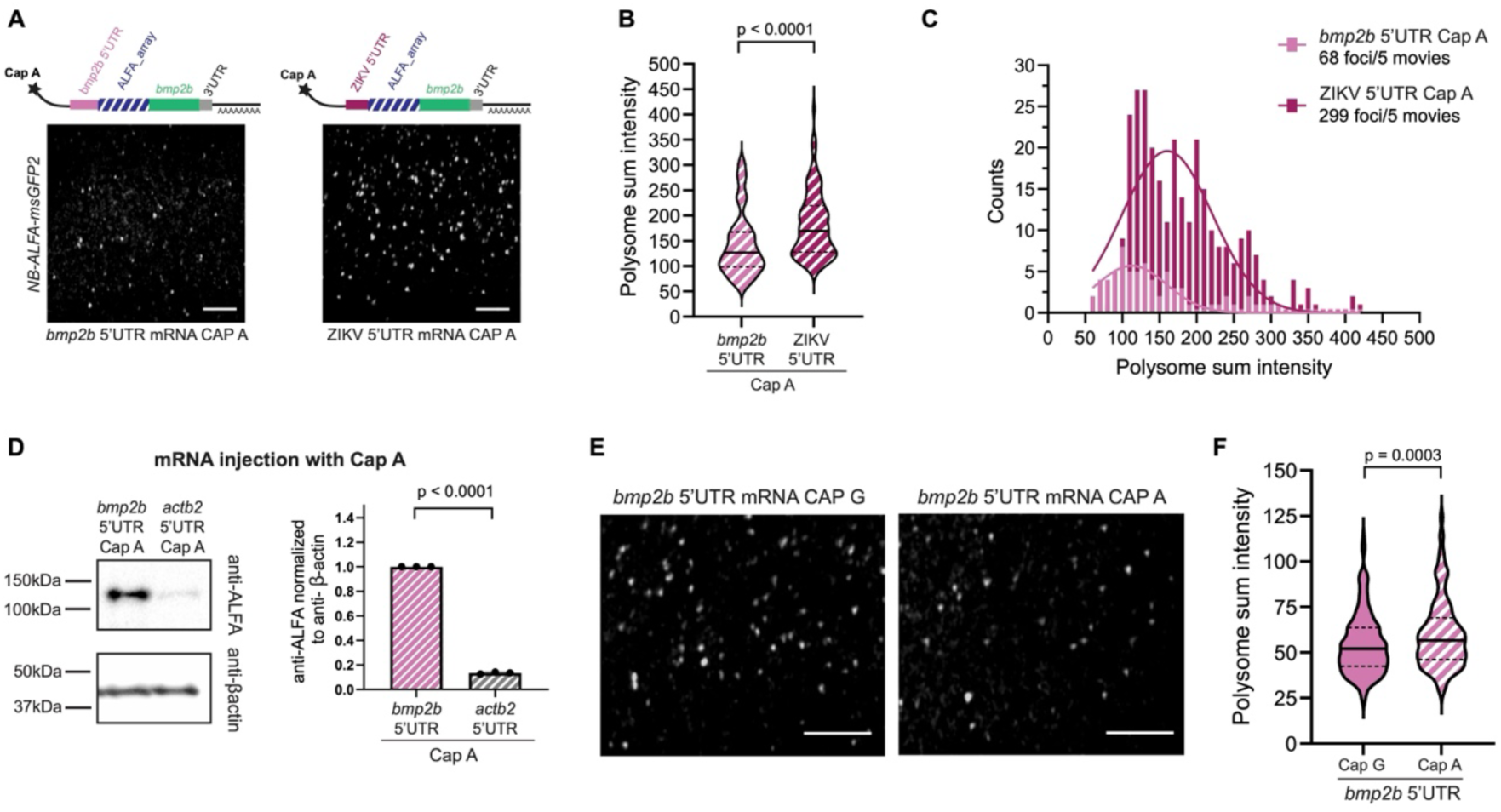
UTR-specific cap-independent translation initiation operates in early zebrafish embryos. (**A**) Schematic of the *in vitro* transcribed mRNA of *ALFA_array_bmp2b* containing the *bmp2b* 5’UTR (pink) or ZIKV 5’UTR (dark pink) with the non-functional cap analog A(ppp)G. Maximum intensity projected images from LLS movies of ∼1.5 hpf *Tg(ubi:NB-ALFA-msGFP2_HIGH)* embryos injected with the respective mRNA. (**B-C**) Violin plot (B) and frequency distribution (C) of polysome sum intensity of *bmp2b* 5’UTR (n= 68 from 5 movies) and ZIKV 5’UTR (n= 299 from 5 movies) *ALFA_array_bmp2b* mRNAs measured from the movies shown in (A). Unpaired *t* test was performed. (**D**) Western blot analysis, using anti-ALFA and anti-β-actin antibodies, of ∼1.5 hpf WT embryos injected with *bmp2b* 5’UTR or *actb2* 5’UTR *ALFA_array_bmp2b* mRNAs with an A(ppp)G (Cap A) cap (left); quantification from three independent crosses and injections (right). Two-tailed Welch’s *t* test was performed. (**E**) Maximum intensity projected images from LLS movies of ∼1.5 hpf *Tg(ubi:NB-ALFA-msGFP2_HIGH)* embryos injected with *bmp2b* mRNAs of *ALFA_array_bmp2b* containing the *bmp2b* 5’UTR and capped with the functional Cap G or non-functional Cap A. (**F**) Violin plot of polysome sum intensity of *ALFA_array_bmp2b* mRNAs containing the *bmp2b* 5’UTR and capped with functional Cap G (n= 301 from 13 movies) or non-functional Cap A (n= 279 from 14 movies) measured from the movies shown in (E). Two-tailed Welch’s *t* test was performed. Scale bars: 10 μm.

## Discussion

In this study, we present a framework for visualizing and quantifying translation dynamics in living vertebrate embryos by implementing the ALFA_array system to zebrafish. Using this approach combined with Lattice Light Sheet microscopy, we were able to directly visualize single *bmp2b* mRNA molecules in a living zebrafish embryo while simultaneously monitoring their translation at single-molecule resolution. Our data indicate that the low translation efficiency of *bmp2b* observed in bulk measurements (*27*, *56*) (fig. S4, D and E), primarily reflects regulation at the level of translation initiation and the limited fraction of mRNAs engaged in translation (Fig. 2E), rather than a reduced elongation rate, which itself falls within the range reported in other species (Fig. 3F) (*69*). Consistent with this, UTR switching with the ZIKV 5’UTR revealed that *bmp2b* translation does not operate at the maximal initiation capacity achievable in early zebrafish embryos (Fig. 4I). This observation suggests that the translational control of *bmp2b* rely mainly on the ability of individual mRNAs to enter translation and initiation rate modulation. We note that because imaging began before reliable embryo orientation (3 hpf) and since *bmp2b* mRNAs become dorsally enriched by 6 hpf, the quantified polysome signals and translation kinetics likely reflect predominantly dorsal translation. While our approach provides key insights into *bmp2b* translation dynamics, future investigation at the endogenous locus will help resolve its regulation within the full physiological context. We note that the N-terminal ALFA-array and associated nanobodies, once bound, may preferentially report cytoplasmic translation. Whether and to what extent Endoplasmic Reticulum-associated translation and subsequent secretory pathway processing contribute to the measured translation kinetics of *bmp2b* remains to be determined.

Removal of the poly(A) tail from the *bmp2b* 3′UTR had only a moderate effect on polysome intensity (Fig. 4B) but reduced protein production (fig. S6A). Therefore, in the context of 2 hpf zebrafish embryos, where poly(A) tail length has been shown to broadly modulate translational efficiency (*62*), the *bmp2b* poly(A) tail appears to support mRNA translatability rather than ribosome loading. In contrast, the *Orthoflavivirus* ZIKV 3′UTR restored stability and enhanced translation, even in the absence of a poly(A) tail, suggesting that it promotes translation initiation through specific structural or sequence elements. These findings support a role for the ZIKV 3′UTR as a poly(A)-independent translation enhancer, consistent with previous *in vitro* observations for other *Orthoflaviviruses* (*70*), but independent of other viral sequences.

Ribosome initiation rate is one of the main limiting factors in translation and it is mostly driven by the 5’UTR (*24*, *25*, *27*). Our data reveal that substitution of the endogenous *bmp2b* 5′UTR with the ZIKV 5′UTR markedly enhanced translation efficiency at the single mRNA level. Strikingly, this enhancement persisted even in the absence of a functional cap, implicating the ZIKV 5′UTR in cap-independent translation, likely *via* an IRES-like mechanism. These data demonstrate *in vivo* that the ZIKV RNA can undergo cap-independent translation, reinforcing prior *in vitro* findings (*29*, *68*), and further show that such mechanisms can operate even without a viral 3′UTR, which is used in most studies (*29*, *68*). Together, our data highlight how viral UTRs can modulate translation, with the ZIKV 5′UTR efficiently hijacking the cellular machinery to maximize ribosome initiation and protein output, thereby enabling the virus to produce proteins as rapidly as possible, and surpassing the translational capacity of typical developmental genes.

In this study, we also show that an endogenous gene, *bmp2b*, is capable of cap-independent translation, uncovering a previously unrecognized mechanism regulating mRNA translation in the early vertebrate embryo. Notably, this mechanism appears to be specific to certain 5′UTRs, including that of *bmp2b*, as we did not observe similar cap-independent translation when testing the 5′UTR of the housekeeping gene *actb2* (Fig. 5D), suggesting that such regulation may be restricted to the 5’UTR of certain developmental genes. While the mechanism remains to be fully elucidated, our findings point to a cap-independent translation enhancer (CITE)-mediated mode of translation (*71–74*), or the presence of an IRES within the potentially highly structured *bmp2b* 5′UTR as suggested by secondary structure predictions (fig. S7, F and G). We found that cap-dependent and cap-independent mechanisms result in similar ribosome engagement in the early zebrafish embryo, consistent with a model in which cap-independent translation is not limited by the initiation rate, but rather by a reduced duration of the translationally active state as predicted in (*75*).

Interestingly, in early zebrafish embryos (i.e., between 1 and 4 hpf), mTOR pathway markers of active canonical translation, including phospho-Rps6 and phospho-eIF4E-BP1, are reduced (*76*). This period of lower canonical translation may provide a window in which non-canonical, cap-independent translation is more permissive, consistent with our observation that *bmp2b* translation can occur efficiently at early embryonic stages.

The early zebrafish embryo thus provides a valuable *in vivo* model to dissect non-canonical translation mechanisms and investigate the contributions of distinct 5′UTR elements to cap-independent gene expression.

In developmental or stress contexts where canonical cap- or poly(A)-dependent translation may be compromised, elements such as IRESs and CITEs can be co-opted to regulate and ensure robust endogenous expression. Non-canonical translation is particularly important in stress responses (*77*) and diseases such as cancer (*78*). Interestingly, cap-independent translation also contributes to stem cell fate decisions *via* DAP5 (eIF4G2/NAT1) (*79*), which mediates the translation of specific genes in human embryonic stem cells (*80*). This non-canonical initiation factor is essential for embryonic development in *Drosophila* (*81*), mouse (*82*), and zebrafish (*83*), thereby highlighting a potentially critical role for cap-independent translation during embryonic development. Whether DAP5 contributes specifically to *bmp2b* translation in zebrafish remains an open question.

Altogether, our findings expand the toolkit for studying gene expression in vertebrates and also reveal how evolutionarily diverse UTRs modulate translation efficiency. Beyond *bmp2b*, whose expression has been well characterized, this platform allows, for the first time, single-molecule resolution analysis of translation in live zebrafish embryos. It opens the door to investigating translational regulation of other developmental genes, providing unprecedented spatiotemporal insights into how translation shapes cell fate, tissue patterning, and morphogen gradients.

## Material and Methods

### Zebrafish Husbandry

Zebrafish husbandry was performed under institutional (MPG) and national (German) ethical and animal welfare regulations. Larvae were raised under standard conditions. Adult zebrafish were maintained in 3.5 Liters tanks at a stock density of 8 zebrafish/L with the following parameters: water temperature: 27 to 27.5 °C; light/dark cycle: 14/10; pH: 7.0 to 7.5; conductivity: 750 to 800 µS/cm. Zebrafish were fed 3 to 5 times a day, depending on age, with granular and live food (*Artemia salina*). Health monitoring was performed at least once a year. All embryos and larvae used in this study were raised at 28.5 °C. All procedures performed on animals conform to the guidelines from Directive 2010/63/EU of the European Parliament on the protection of animals used for scientific purposes and were approved by the Animal Protection Committee (Tierschutzkommission) of the Regierungspräsidium Darmstadt (reference: B2/1218).

Zebrafish housing and husbandry was approved by the “Comité d’Etique pour l’Expérimentation Animale et la Direction Sanitaire et Vétérinaire de l’Hérault” (Aquatic model facility ZEFIX CRBM, C-34-172-39).

### Animal Experimentation

All experiments involving zebrafish were carried out on embryos/larvae before 5 dpf and complied with the guidelines of the European Parliament Directive 2010/63/EU on the protection of animals used for scientific purposes.

### Plasmid generation for zebrafish line establishment

The plasmids used to generate the *Tg(ubi:NB-ALFA-msGFP2_HIGH)bns698*, *Tg(ubi:NB-ALFA-msGFP2_LOW)bns699*, *Tg(ubi:scFv-msGFP2)bns715*, *Tg(ubi:NB-mScarlet3)bns740* lines, were generated using NEBuilder® HiFi DNA Assembly from published ubiquitin promoter (*45*), NB-msGFP2-GB1-NLS (*38*), scFv-msGFP2-GB1-NLS (*38*) sequences in Tol2 plasmid containing eCFP under the control of gamma-crystallin promoter (gift from Darren Gilmour, Department of Molecular Life Sciences, University of Zurich). All the three nanobodies were fused to a GB1 (B1 domain of streptococcal protein G) to improve solubility and a Nuclear Localization Signal (NLS) to the scFv and NB-msGFP2 to improve the signal-to-noise ratio in the cytoplasm. For the *Tg(bmp2b:ALFA_array_bmp2b)bns739, bmp2b* promoter (primer sequences from (*16*)), UTRs and coding sequence are from WT(AB) genomic DNA. The *ALFA_array* sequences from (*38*) were codon optimized for zebrafish (https://en.vectorbuilder.com/tool/codon-optimization.html) and synthetized at GenScript Biotech Corporation. The final plasmid was generated using NEBuilder® HiFi DNA Assembly.

Transgenesis was done by injection of one-cell stage embryos with 20 pg of plasmid DNA together with 15 pg of transposase *Tol2* mRNA and 0,2% phenol red. The positive offspring were outcrossed with WT(AB) for 2 generations before establishing the stable line.

Full sequences obtained with whole plasmid sequencing are available in Supplementary Table 1.

### Plasmid generation for *in vitro* transcription

Bmp2b UTRs and coding sequence were amplified from cDNA since it contains introns, the *ALFA_array* sequences were from (*38*) and a T7 sequence (TAATACGACTCACTATAGGG) was added and assembled using NEBuilder® HiFi DNA Assembly. From this plasmid, 5’ or 3’UTR of *bmp2b* were removed and 5’ (5’-AGTTGTTGATCTGTGTGAATCAGACTGCGACAGTTCGAGTTTGAAGCGAAAGCTAGCAACAGTATCAACAGGTTTTATTTTGGATTTGGAAACGA GAGTTTCTGGTC-3’) or 3’UTR (5’-GCACCAATCTTAATGTTGTCAGGCCTGCTAGTCAGCCACAGCTTGGGGAA AGCTGTGCAGCCTGTGACCCCCCCAGGAGAAGCTGGGAAACCAAGCCTATAGTC AGGCCGAGAACGCCATGGCACGGAAGAAGCCATGCTGCCTGTGAGCCCCTCAGA GGACACTGAGTCAAAAAACCCCACGCGCTTGGAGGCGCAGGATGGGAAAAGAA GGTGGCGACCTTCCCCACCCTTCAATCTGGGGCCTGAACTGGAGATCAGCTGTGG ATCTCCAGAAGAGGGACTAGTGGTTAGAGGAGACCCCCCGGAAAACGCAAAACA GCATATTGACGCTGGGAAAGACCAGAGACTCCATGAGTTTCCACCACGCTGGCCG CCAGGCACAGATCGCCGAATAGCGGCGGCCGGTGTGGGGAAATCCATGGGTCT-3’) of Zika were amplified from a molecular clone of the BeH819015 isolate representative of the most prominent group of ZIKV isolates from South America as it differs very few from other sequenced isolates (*84*) and assembled using NEBuilder® HiFi DNA Assembly. The 30 MS2 loops were reengineered from (*85*) using prediction with (*86*) to obtain 30x functional, short and low affinity MS2 sequences and synthetized at GenScript Biotech Corporation, and were added after the stop codon of *bmp2b*.

Full sequences obtained with whole plasmid sequencing are available in Supplementary Table 1.

### *In vitro* transcription and injections of mRNAs

Capped mRNAs were synthesized using the HiScribe® T7 ARCA mRNA Kit (New England Biolabs, #E2065) following the manufacturer protocol. Poly(A) tailing was perfomed using E. coli Poly(A) Polymerase kit (New England Biolabs, M0276). Purification of mRNAs were done with Zymo Research RNA Clean & Concentrator kit (R1016). For Cap analog mRNA synthesis, MEGAshortscript™ T7 Transcription kit (Invitrogen, AM1354) was used following the manufacturer protocol and adding the unmethylated G(5’)ppp(5’)A RNA Cap Structure Analog (New England Biolabs, #S1406) or 3′-O-Me-m7G(5’)ppp(5’)G (New England Biolabs, #S1411) in excess (2 mM each NTP and 8 mM Cap analog).

For western blot analysis, one-cell stage embryos were injected with 25 pg of mRNA in 1 nl volume of water. For Lattice Light sheet imaging, one-cell stage embryos were injected with 2.5 pg of mRNA in 1 nl volume of water.

For puromycin experiment, 25 pg of mRNA in 1 nl volume of water with or without 10mg/mL concentration of puromycin (Merck, P8833) were injected.

### Plasmid generation for transient experiments

*ALFA_array_bmp2b* plasmid was used from the generation of the transgenic line, and Zika virus 5’UTR (see section plasmid generation for *in-vitro* transcription for the sequence) was added using NEBuilder® HiFi DNA Assembly.

For transient experiments, 20 pg of plasmids were injected without phenol red. All reagents were injected at a volume of 1 nl.

### Immuno-HCR-FISH

Immunostaining coupled with Hybridization Chain Reaction Fluorescence In-Situ Hybridization (HCR-FISH, Molecular Instruments®) was performed as described in (*87*) and according to manufacturer’s protocol with few modifications. Embryos were fixed at the desired stage in 4% paraformaldehyde (PFA) for 16 hours at 4°C. Then embryos were washed 3 times 5 min with 1mL phosphate buffer saline (PBS). At that point, embryos were manually dechorionated using forceps. Embryos were incubated 4 times 10 min in methanol (MeOH) and once for 50 min. Embryos were stored at least overnight at −20°C in MeOH. Then, ∼9 embryos per tube were rehydrated with series of MeOH/1XPBS-0.1%Tween-20 (PBST) washes for 5min at room temperature (75% MeOH, 50% MeOH, 25% MeOH and 5 times PBST). Embryos were pre-hybridized with 250 μL of hybridization solution (Molecular Instruments®) for 30 min at 37°C and hybridized with 16 nM of probe in 250 μl of hybridization solution overnight at 37°C. Probes were designed by Molecular Instruments® by giving the sequence of the ALFA_array (see Supplementary Table 1). The next day, embryos were washed 3 times 15 min with 250 μl wash buffer (Molecular Instruments®) at 37°C followed by 2 times 5 min in 500 μl 5× SSC 0.1%Tween-20 (SSCT) at room temperature. Amplification step was done with a 30 min pre-amplification wash in 250μl amplification buffer (Molecular Instruments®) followed by incubation in 250μl amplification buffer with 30pmol of each hairpin (h1 and h2, HCR™ Amplifier (v3.0) 647 dye) overnight in the dark. Then embryos were washed 2 times 5 min, 2 times 30 min and once 5 min in 5× SSCT. Then the samples were washed 3 times 5 min in PBST and blocked with 0.5% ultrapure BSA (Invitrogen™ AM2616) in PBST for 2 hours at room temperature. Embryos were then incubated overnight at 4 °C with rabbit anti-ALFA (1:1000, N1581 Nanotag Biotechnologies) in block solution. The next day embryos were washed 6 times 30min in PBST and incubated with 1/500e secondary antibody (goat anti-rabbit Alexa Fluor™ 568 A11036 ThermoFisher) overnight at 4 °C. Then embryos were washed 6 times 30min in PBST and mounted in ProLong™ Diamond with DAPI (Invitrogen P36962). Embryos were imaged on a Zeiss LSM800 Observer Airyscan.

### Whole-mount fluorescent *in situ* hybridization

The in situ probe for *ALFA_array* (of 800 nucleotides length) was synthesized from a PCR product from the *bmp2b:ALFA_array_bmp2b* plasmid using the following primers: *ALFA_array*-ISH forward: 5′-GAAGCGGAAGCGGACCATCTAG-3′ and *ALFA_array*-ISH reverse: 5′ CAGATCAATTAACCCTCACTAAAGGGAGAAGGTCCACTTCCAGATC-3′.T3 RNA polymerase (Roche, Cat. No. 11031163001) was used for transcription of probe using digoxigenin-labeled NTPs (Roche, Cat. No. 11277073910). Adult *Tg(bmp2b:ALFA_array_bmp2b)* fish were intercrossed and embryos were fixed at the desired time points in 4% PFA overnight at 4 °C. Then, embryos were dehydrated through a graded MeOH series, with each step lasting 5 min: 25% MeOH, 50% MeOH, 75% MeOH, and 100% MeOH. They were subsequently stored at −20 °C in MeOH until needed. On the first day, after rehydration and washing in PBS-0.1% Tween-20 (PBST), embryos were prehybridized in hybridization buffer (50% formamide, 5× SSC, 0.1% Tween-20, 9.2 mM citric acid, 50 μg/mL heparin, 500 μg/mL of tRNA) at 70 °C for 2 hours, then incubated overnight at 70 °C with 1 ng/μL DIG-labeled RNA probe in hybridization buffer. On the second day, embryos were subjected to a series of washes at 70 °C, transitioning from 100% hybridization buffer to 2× SSC in 10-minute steps (25%, 50%, 75% and 100% 2× SSC). This was followed by room-temperature washes in 75%, 50%, and 25% 0.2× SSC in maleic acid buffer (100mM maleic acid and 150mM NaCl, pH 7.5) containing 0.1% Tween-20 (MABT) for 10 min each, with a final 10 min wash in MABT. Embryos were then incubated in 2% blocking reagent (Roche, Cat. No. 11096176001) in MABT for 1 hour, followed by overnight incubation at 4 °C with a 1:2000 dilution of anti-digoxigenin-POD antibody (Roche, Cat. No. 11207733910) in blocking buffer. On the third day, after additional washes (6 times, 10 min in PBST), embryos were stained using a 1:100 dilution of tyramide-Fluorescein reagent in amplification buffer (Akoya Biosciences, NEL741001KT) for 30 min at room temperature. After washing in PBST for 3 times 10 min each, embryos were stain with DAPI and mounted in ProLong™ Diamond. Imaging was performed on a ZEISS Celldiscoverer 7 automated microscope equipped with an LSM 900 confocal module and Airyscan 2 detector using a Plan-APOCHROMAT 20X / 0.95NA W objective.

### Light Sheet live imaging

Embryos were collected after laying and mounted in 0.75% low-melting agarose (Sigma-Aldrich®, A4018) in a capillary immerged with egg water (Instant Ocean® Aquarium Systems, with miliQ water). Embryos were imaged on a Light Sheet Z.1 with the following settings: 20x/1.0 Corr DIC M27 water objective, images (1920 × 1920 pixels) for msGFP2 a 488nm laser was used and 160z-positions were taken with 1µm z-step (37.8 seconds per frame). For *Tg(ubi:NB-ALFA-msGFP2_HIGH)bns698* and *Tg(ubi:NB-ALFA-msGFP2_LOW)bns699* embryos with zygotic-only expression, images were acquired at 4 minute intervals to accommodate longer recording durations and minimize data volume. Movies were maximum intensity projected and processed with Fiji (*88*).

### Fluorescence recovery after photobleaching

Fluorescence recovery after photobleaching (FRAP) was performed in zebrafish embryos expressing NB-ALFA-msGFP2 which were injected with 20pg of H2B_mCherry_ALFAtag plasmid (*51*) and imaged 1 hour later at 28.5 °C. The following settings were used: a Zeiss LSM880 using a 40x/1.3 Oil objective. Images (512 × 512 pixels), 16 bits/pixel, pixel size 0.104 µm, zoom 4x, were acquired every ∼6 sec during 43 frames. Excitation: GFP with a laser at 488nm and mCherry with a laser at 561 nm. Laser power was measured using the 10×/0.45 Plan-Apochromat Air objective and was ∼1.2 mW for 488 nm and ∼4.2 mW for 561 nm at 100% laser. For acquisition, measured laser power was ∼60 µW (488 nm) and ∼83 µW (561 nm). A circular ROI (3 µm diameter) was bleached using hundred pulses of 488 nm and 561 lasers at maximal power (100%) after 4 pre-bleach frames. To discard any source of fluorescence intensity recovery other than molecular diffusion, the measured fluorescence recovery in the bleached ROI region (*I_bl_*) was corrected by an unbleached ROI (*I_unbl_*) of the same nucleus and another ROI outside of the nuclei (*I_out_*) following the simple equation:

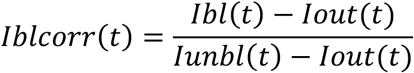

### Confocal live imaging

Confocal live imaging was performed on a Zeiss LSM880 Airyscan with 40x/1.1 water objective. For NB-msGFP2 and scFv-msGFP2 imaging, we used the following settings: images (512 x 512 pixels), 488 nm laser, zoom x 2, 30 z-positions were taken with 1.5 µm z-step and a pixel size of 0.28 µm (43.8 seconds per frame).

### Lattice Light Sheet live imaging

After injection, embryos were manually dechorionated using forceps and mounted in 0.75% low-melting agarose (Sigma-Aldrich®, A4018) with egg water in a Fluorodish (Dutscher FD35-100). Embryos were imaged on a Lattice Light Sheet 7 (Zeiss LLS7) with the following settings:

For mRNA molecules imaging of *Tg(actb2:2xMCP-NLS-EGFP)* embryos injected with *ALFA_array_bmp2b_30XMS2*: 488 nm laser was used with 20 ms exposure time and 2x-plane were taken with 0.5 μm x-step for 200 time series, image size are 1024 x 512 pixels (180 milisecond per frame).

For simultaneous mRNA and translation imaging of *Tg(ubi:NB-ALFA-mScarlet3); Tg(actb2:2xMCP-NLS-EGFP)* embryos injected with *ALFA_array_bmp2b_30XMS2* : 488 nm laser was used with 30ms exposure time and 561nm laser was used with 30ms exposure time and 70x-plane were taken with 0.5 μm x-step for 10 time series, image size are 478 x 2048 pixels (943 ms per frame).

For kinetics measurements of *Tg(bmp2b:ALFA_array_bmp2b)bns739*: a 488nm laser was used at ∼265 μW (measured after the objective using a Thorlabs laser power meter). Images were acquired with an exposure time of 10 ms. A total of 70 x-planes were collected with a x-step of 0.5 µm for each of the 180 time points. The image size was 674 × 2048 pixels, and corresponds to an acquisition time of 1.19 sec per frame.

For kinetics measurements of *Tg(ubi:NB-ALFA-msGFP2_HIGH)* embryos injected with *bmp2b* 5’UTR DNA or ZIKV 5’UTR DNA: a 488 nm laser was used at ∼265 μW (measured after the objective using a Thorlabs laser power meter). Images were acquired with an exposure time of 10 ms. A total of 70 x-planes were collected with a x-step of 0.5 µm for each of the 180 time points. The image size was 674 × 2048 pixels, and corresponds to an acquisition time of 1.19 sec per frame.

For mRNA injections with/without puromycin: a 488 nm laser was used at ∼305 μW (measured after the objective using a Thorlabs laser power meter) with 10ms exposure and single x-plane, image size are 674 x 2048 pixels.

For spot intensity analysis: a 488nm laser was used at ∼305 μW (measured after the objective using a Thorlabs laser power meter) with 10 ms exposure time and 70x-plane were taken with 0.5 μm x-step for 30 time series, image size are 674 x 2048 pixels.

Images were processed with the Zeiss Lattice Light Sheet processing tool using “Deskew>Cover Glass Transfomation” tool from Zen Blue (2.3 lite) software.

### Kinetics measurement analysis

Movies for extraction of translation kinetics from *Tg(bmp2b:ALFA_array_bmp2b)bns739* or embryos injected with *bmp2b* 5’UTR DNA or ZIKV 5’UTR DNA, were acquired on Lattice Light Sheet (Zeiss LLS7) and were cover glass transformed and maximum intensity projected with Zen Blue (2.3 lite) software. Fluorescence intensity fluctuations of translation sites were obtained using spot detector Fiji (*88*) plugin TrackMate LOG (Laplacian Of Gaussian) and tracking was done with the TrackMate Linear Assignment Problem (LAP) from Fiji plug-ins (*89*, *90*). In order to generate the tracked intensity fluctuation, each detected spot was divided by background at each time frame. Tracks were then selected if their time length was superior to 150 frames (178.5 sec).

The multiple tau algorithm allows to avoid overweighting of the end of the correlation versus the beginning.

The elongation dwell time T was obtained by fitting the measured auto-correlation, with the following analytical expression as in (*33*, *34*, *37*):

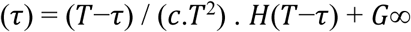

where *c* is the translation initiation rate and *H*(*t*) is the Heaviside step function equal to 1 for *t*>0 and zero otherwise. We included a G∞ term in the fit as in (*37*), to account for a non-zero baseline at long correlation times, which can arise from instrumental offsets, background noise, or slow signal drifts. This prevents bias in the estimation of the correlation amplitude and the derived parameters such as the particle number. Autocorrelation curves were fitted with a Levenberg-Marquadt algorithm using Python.

### Polysome intensity measurements

Polysome intensity measurement was performed using ICY software (*91*). Movie files were imported into ICY, and the Spot Detector plugin was used with a manually defined threshold to identify objects. Only polysome intensities from the first frame were retained (to avoid photobleaching bias) for analysis. Background intensity was measured by selecting regions of interest (ROIs) devoid of translation dots. For each movie, the mean background intensity was calculated from at least five ROIs. Finally, each spot was divided by the respective value of the background. All graphs were generated using GraphPad Prism 10.2.3 (403). Outliers were removed using ROUT method in GraphPad Prism with lowest stringency (Q=0.1%). Frequency distribution was done with a binning of 10. For statistical tests, first a *F*-test of equality of variances was performed. If the variances were equal, two-tailed unpaired *t* test was performed. If the variance were not equal, a two-tailed Welch’s *t* test was performed. In the violin plots, plain black lines represent the median and the dashed lines the quartiles.

### Western blot analysis

Embryos were collected at the indicated times post-fertilization. For 1.5, 3, 4.3 and 6 hpf embryos the yolks were removed by hand with a scalpel. For 9, 12 and 24 hpf embryos, the yolks were removed using deyolking buffer (55mM NaCl, 1.8 mM KCl, 1.25 mM NaHCO_3_). Embryos were then snap frozen in liquid nitrogen. Embryos were crushed with a pestle with 4uL per embryo of 4x Laemmli Sample Buffer (Bio-Rad 1610747) supplemented with 2-mercaptoethanol. Samples were then heated at 70°C for 10 min, the volume equivalent to 2,5 embryos per well was loaded on a 4-12% Bis-Tris NuPAGE™ gel (Invitrogen NP0335), and ran at 180 V for 1 h. Protein transfer was done at 4°C for 1 h at 110 V onto a 0.2 μm nitrocellulose membrane (Invitrogen LC2000). The membrane was blocked in 5% milk-PBT (PBS+0.1% Tween 20) for 40 min and incubated overnight at 4 °C with rabbit anti-ALFA (1:1000, N1581 Nanotag Biotechnologies) or mouse anti-β-actin (1:1000, Sigma A5441) primary antibodies in PBT. Anti-mouse (1:5000, Abcam ab97023) and anti-rabbit (1:5000, Cell Signaling 7074) HRP-linked secondary antibodies were used and incubated 1 h at room temperature in 5% milk-PBT. Chemiluminescent detection was performed using the SuperSignal™ West Pico PLUS chemiluminescent substrate kit (ThermoFisher 34577) on a Biorad ChemiDoc™ MP Imaging System. Signal quantification of western blots was perfomed using Fiji (*88*) (mean intensity was taken for each band).

### Mean square displacement analysis

Movies were first coverglass transformed and maximum intensity projected with Zen software, and single particles were detected using spot detector Fiji plugin TrackMate LOG (Laplacian Of Gaussian) and tracking was done with the TrackMate Linear Assignment Problem (LAP) (*89*, *90*). In total n=2729 traces, from 8 movies for *bmp2b* 5’UTR construct and n=3464 traces from 13 movies for ZIKV 5’UTR construct. 1D displacement in x and y were obtained from TrackMate files from Fiji for each movie. Mean square displacements (MSD) were calculated for tracks present for at least 20 consecutive frames using the MSDanalyzer MatLab script (*92*) on Matlab_R2024b software version. Instantaneous coefficients diffusion were obtained by fitting 0 to 20 frames of the MSD of individual trajectories with a linear model as follows:

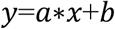

To estimate MSD, according to Einstein’s theory, the MSD of Brownian motion is described as:

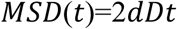

with d representing dimensionality, in this case, d = 2. D represents the diffusion coefficient. A linear fitting was considered unsatisfactory when R^2^<0.8 and was removed from the dataset.

## Acknowledgements

We thank Dr. Nadine Vastenhouw for providing the *Tg(actb2:2xMCP-NLS-EGFP)* line, Oliver Buchhold and Rita Retzloff from the Max Planck Institute for Heart and Lung Research fish facility and Aurélien Drouard, Marc Plays, Anthony Ollier, Anne Morel, and Bénédicte Delaval from the Aquatic model platform Biocampus ZEFIX, for technical advice and support, Dr. Simon Perathoner for administration and comments on the manuscript, Dr. Mario Looso for help with RNAseq analysis, the Montpellier Resource Imaging (MRI) platform, in particular Dr. Virginie Georget, for their support with the Zeiss Lattice Light Sheet 7 microscope, and Dr. Frank Martin for his valuable feedback.

## Author Contributions

MB conceived the study, with support from JD for the viral work and from DYRS for the zebrafish work. MB designed the methods with the help of JD. MB, JD, JL, ML, DA, HM and VM performed the experiments. KM and MB analyzed the imaging data, and TM performed the data fitting. DM, TJS, and DYRS provided resources. MB wrote the first draft of the manuscript, and all authors contributed to its revision and approved the final version. MB and DYRS coordinated project administration and funding.

## Data, Code, and Materials Availability

All the data and code needed to evaluate and reproduce the results in the paper are present in the paper and/or the Supplementary Materials. Materials created in this study (e.g., fish lines, plasmids) are available upon request to the corresponding authors.

