## Supplementary figures and table1 for "Real-time single-molecule imaging in zebrafish embryos uncovers non-canonical translation"

Maëlle Bellec *et al.*

**This PDF file includes:**

Figs. S1 to S7
Table S1
Movie legends S1 to S6

**Other Supplementary Materials for this manuscript include the following:**

Movies S1 to S6

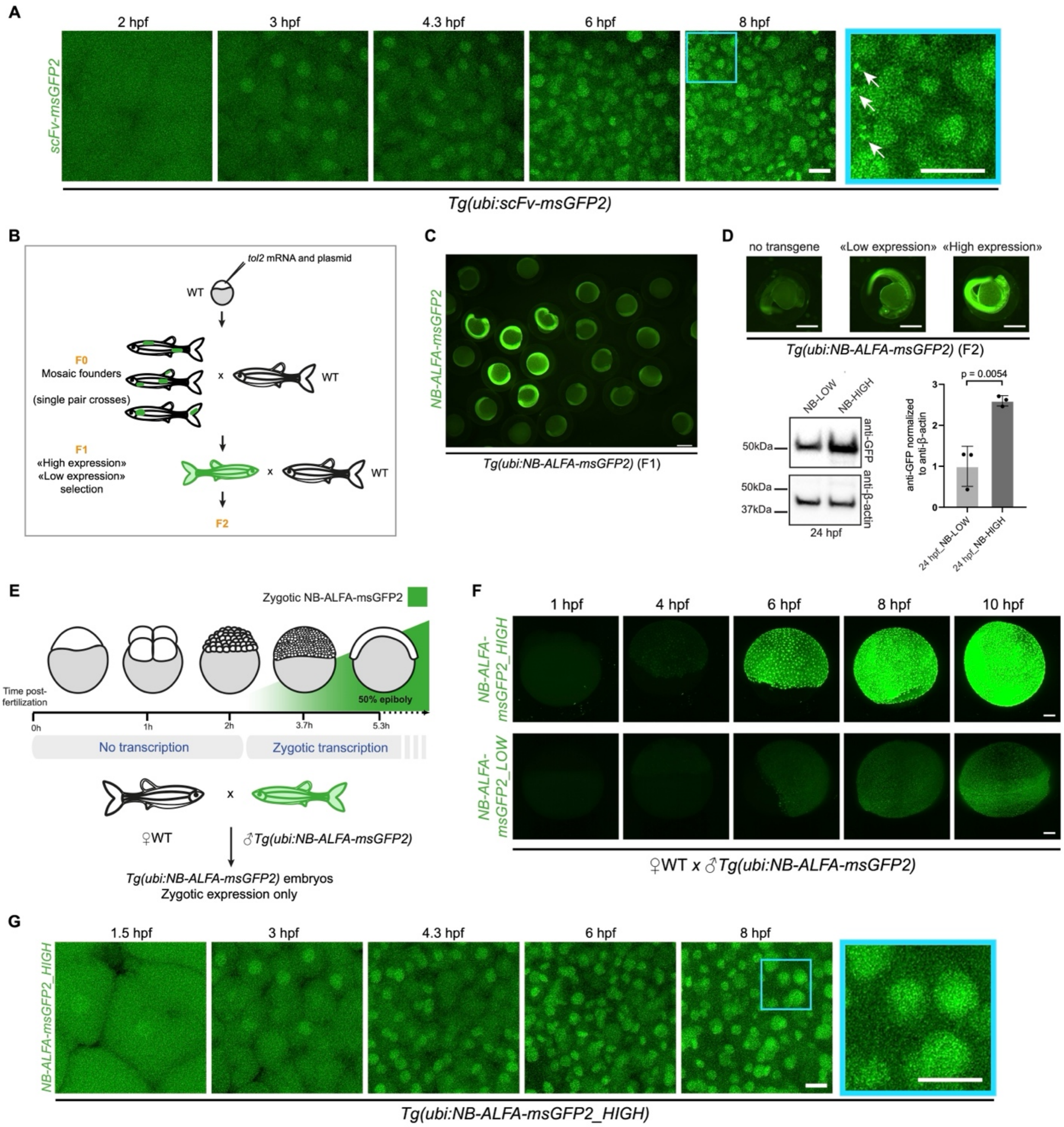


**fig. S1. Zygotic expression of the NB-ALFA-msGFP2 transgene and absence of aggregates compared with the scFv-msGFP2 transgene.** (**A**) Maximum intensity projected images from a confocal movie of a *Tg(ubi:scFv-msGFP2)* embryo. (**B**) Schematic of the crosses used to generate the transgenic lines expressing NB-ALFA-msGFP2. (**C**) Wide field image of F1 embryos expressing different levels of NB-msGFP2. (**D**) Wide field images of 24 hpf embryos without a transgene, or with low and high expression of NB-ALFA-msGFP2. Western blot analysis using anti-GFP and anti-β-actin antibodies of 24 hpf *Tg(ubi:NB-ALFA-msGFP2_LOW)* and *Tg(ubi:NB-ALFA-msGFP2_HIGH)* embryos; quantification from three replicates, error bars are standard deviation. Unpaired *t* test was performed. (**E**) Schematic representing the maternal-to-zygotic transition in zebrafish with the zygotic NB-ALFA-msGFP2 expression in green. Below, schematic of the cross used to study zygotic expression of NB-ALFA-msGFP2. (**F**) Maximum intensity projected images of light sheet movies of *Tg(ubi:NB-ALFA-msGFP2_HIGH)* and *Tg(ubi:NB-ALFA-*

*msGFP2_LOW)* embryos expressing the anti-ALFA-msGFP2 nanobody zygotically. (**G**) Maximum intensity projected images from a confocal movie of embryo expressing NB-ALFA-msGFP2_HIGH. Scale bars: 20 µm (A, G), 500 µm (C, D), 100 µm (F).

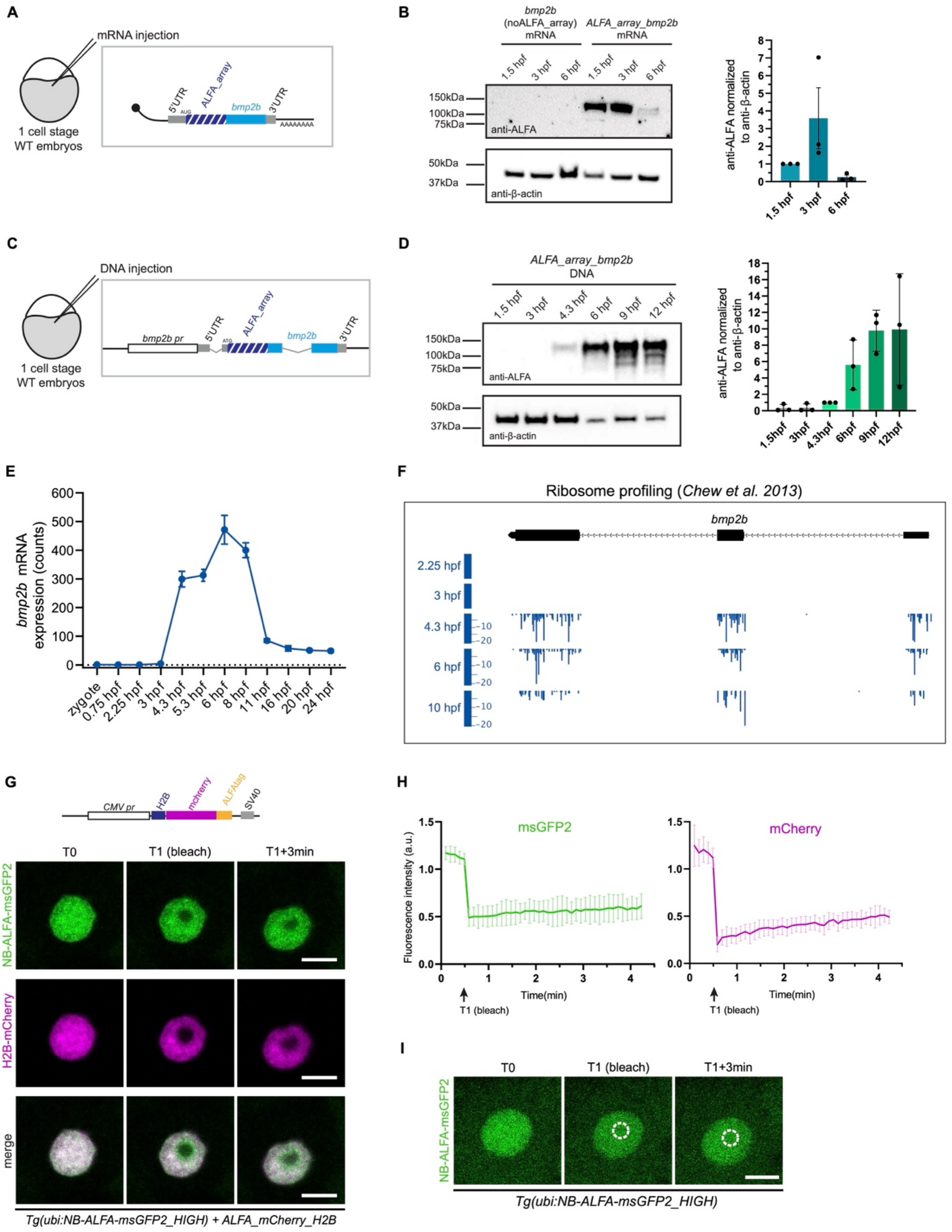


**fig. S2. *bmp2b* time course expression and stable binding affinity of the anti-ALFA nanobody to the ALFA-tag in zebrafish embryos.** (**A**) Schematic of the mRNA containing *bmp2b* fused to the ALFA_array tag injected into one-cell stage WT embryos. (**B**) Western blot analysis, using anti-ALFA and anti-β-actin antibodies, of WT embryos injected with *ALFA_array_bmp2b* mRNA, and collected at the indicated stage; quantification from three biological replicates. (**C**) Schematic of the plasmid containing *bmp2b* fused to the ALFA_array tag injected into one-cell stage WT embryos. (**D**) Western blot analysis, using anti-ALFA and anti-β-actin antibodies, of WT embryos injected with *ALFA_array_bmp2b* DNA, and collected at the indicated stage; quantification from three biological replicates. (**E**) Graph representing *bmp2b* mRNA expression at different time points from RNA sequencing data (*47*) (**F**) Ribosome profiling data of *bmp2b* mRNA at 2.25, 3, 4.3, 6 and 10 hpf from (*48*). (**G**) On top, schematic of the *H2B_mCherry_ALFA-tag* plasmid used for the FRAP experiments from (*51*). On the bottom, images from a FRAP experiment on a nucleus of a ~5 hpf *Tg(ubi:NB-ALFA-msGFP2_HIGH)* zebrafish embryo (NB-ALFA-msGFP2 in green) injected with *H2B_mCherry_ALFA-tag* DNA (magenta). Images before and after photobleaching are shown. (**H**) FRAP recovery curve from experiment in (G) of the fluorescence intensity as a function of time. Recovery of NB-ALFA-msGFP2 fluorescence is in green and of H2B_mCherry fluorescence in magenta (n=7, from three embryos). (**I**) Images from a FRAP experiment on a nucleus of a *Tg(ubi:NB-ALFA-msGFP2_HIGH)* zebrafish embryo (NB-ALFA-msGFP2 in green) but without *H2B_mCherry_ALFA-tag*. Images before and after photobleaching are shown. The bleached area is encircled by a dashed white line. Scale bars: 10 µm (F, H).

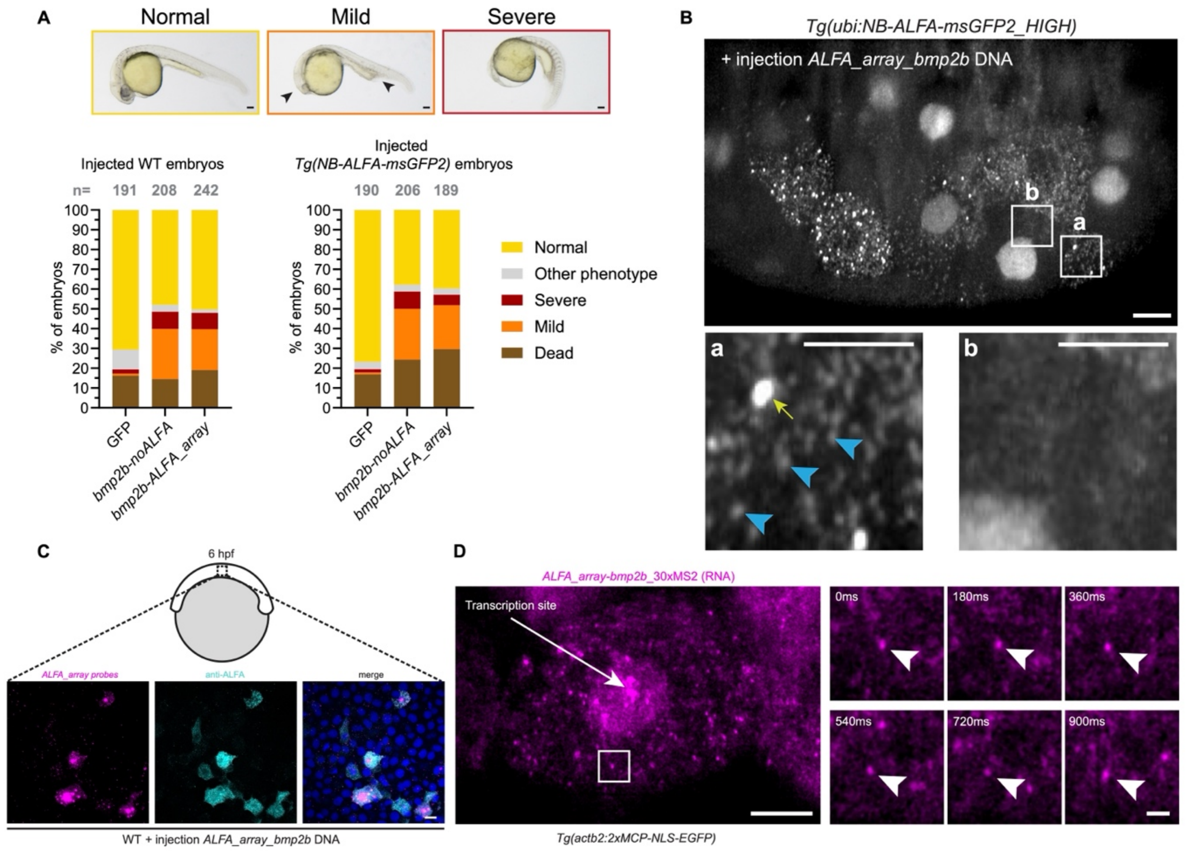


**fig. S3. Lattice Light Sheet imaging of nascent *bmp2b* translation and mRNA molecules in zebrafish embryos.** (**A**) Quantification of ventralization phenotypes at 24 hpf upon injection into WT embryos of the *ALFA_array_bmp2b* mRNA*, bmp2b* without the ALFA_array, or GFP mRNA. Upper panels show representative images of embryos with the respective phenotype. Arrowheads point to a reduced head size and enlarged ventral fin fold. (**B**) Maximum intensity projected image from an LLS movie of a ~4.3 hpf *Tg(ubi:NB-ALFA-msGFP2_HIGH)* embryo injected with *ALFA_array_bmp2b* DNA. Zoomed in panels show cells with (a) or without (b) ALFA_array_Bmp2b expression. Yellow arrow points to a translation dot. Blue arrowheads point to putative single molecules of ALFA-tagged Bmp2b protein. (**C**) Maximum intensity projection of confocal images from immuno-HCR-FISH with *ALFA_array* probes (magenta) and anti-ALFA antibody (cyan) on a 6 hpf WT embryo injected with *ALFA_array_bmp2b* DNA. (**D**) Images from an LLS movie of a ~4.3 hpf *Tg(actb2:2xMCP-NLS-EGFP)* embryo injected with *ALFA_array_bmp2b_30xMS2* DNA. MCP-NLS-EGFP is represented in magenta and corresponds to mRNA molecules. Larger view is shown on the left with an arrow pointing to a transcription site and zoomed in panels on the right with arrowheads pointing to one mRNA molecule in motion. Scale bars: 100 µm (A), 10 µm (B, C, D), 1 µm (zoomed in panels B, D).

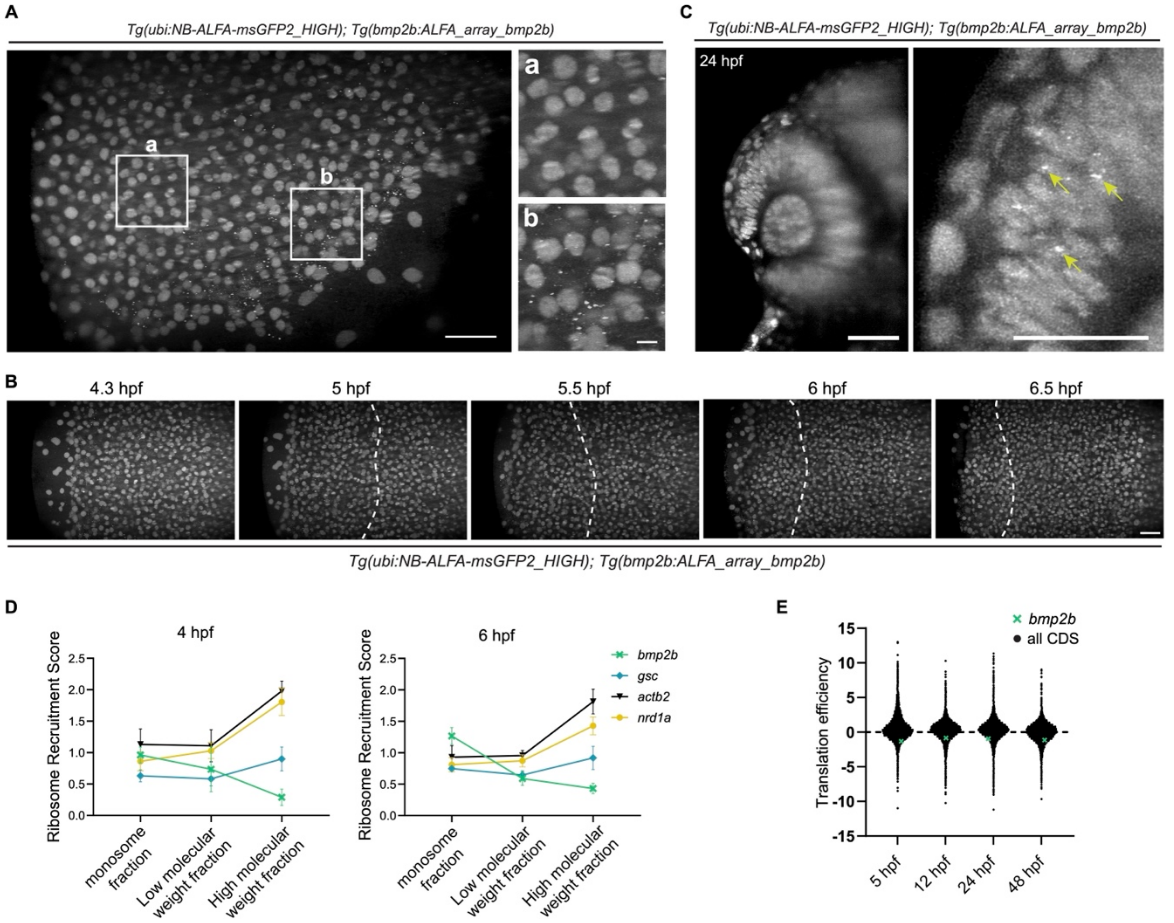


**fig. S4. Developmental pattern and efficiency of *bmp2b* translation.** (**A**) Maximum intensity projected images from an LLS movie of a ~4 hpf *Tg(ubi:NB-ALFA-msGFP2_HIGH); Tg(bmp2b:ALFA_array_bmp2b)* embryo. Yolk is on the bottom right of the image. Zoomed in images outside (a) and inside (b) *bmp2b* expression pattern are on the right. (**B**) Maximum intensity projected images from an LLS movie of a *Tg(ubi:NB-ALFA-msGFP2_HIGH); Tg(bmp2b:ALFA_array_bmp2b)* embryo from ~4.3 to ~6.5 hpf. Dashed white lines delineate the border of the *bmp2b* translation zone which is to the left of the line. The translation foci progressively shift toward the embryo's periphery as development proceeds. Animal pole is on the right. (**C**) Maximum intensity projected images from an LLS movie of a 24 hpf *Tg(ubi:NB-ALFA-msGFP2_HIGH); Tg(bmp2b:ALFA_array_bmp2b)* embryo. A zoomed in image of the dorsal retina is on the right. Yellow arrows point to translation dots. (**D**) Ribosome recruitment score (the ratio between reporter abundance in the ribosome-bound fraction and the total RNA pool) for several genes (5'UTRs upstream of a reporter) at 4 and 6 hpf, *bmp2b* reads are mostly present in the monosome fraction. Data from (*27*). (**E**) Translation efficiency (calculated by dividing ribosome protected fragment levels (in Reads Per Kilobase per Million (RPKM) by mRNA expression (in RPKM)) for all coding sequences (CDS) at the indicated timepoints. *bmp2b* is highlighted in green. Data from (*56*). Scale bars: 50 µm (A, B, C), 20 µm (zoomed in panel C), 10 µm (zoomed in panel A).

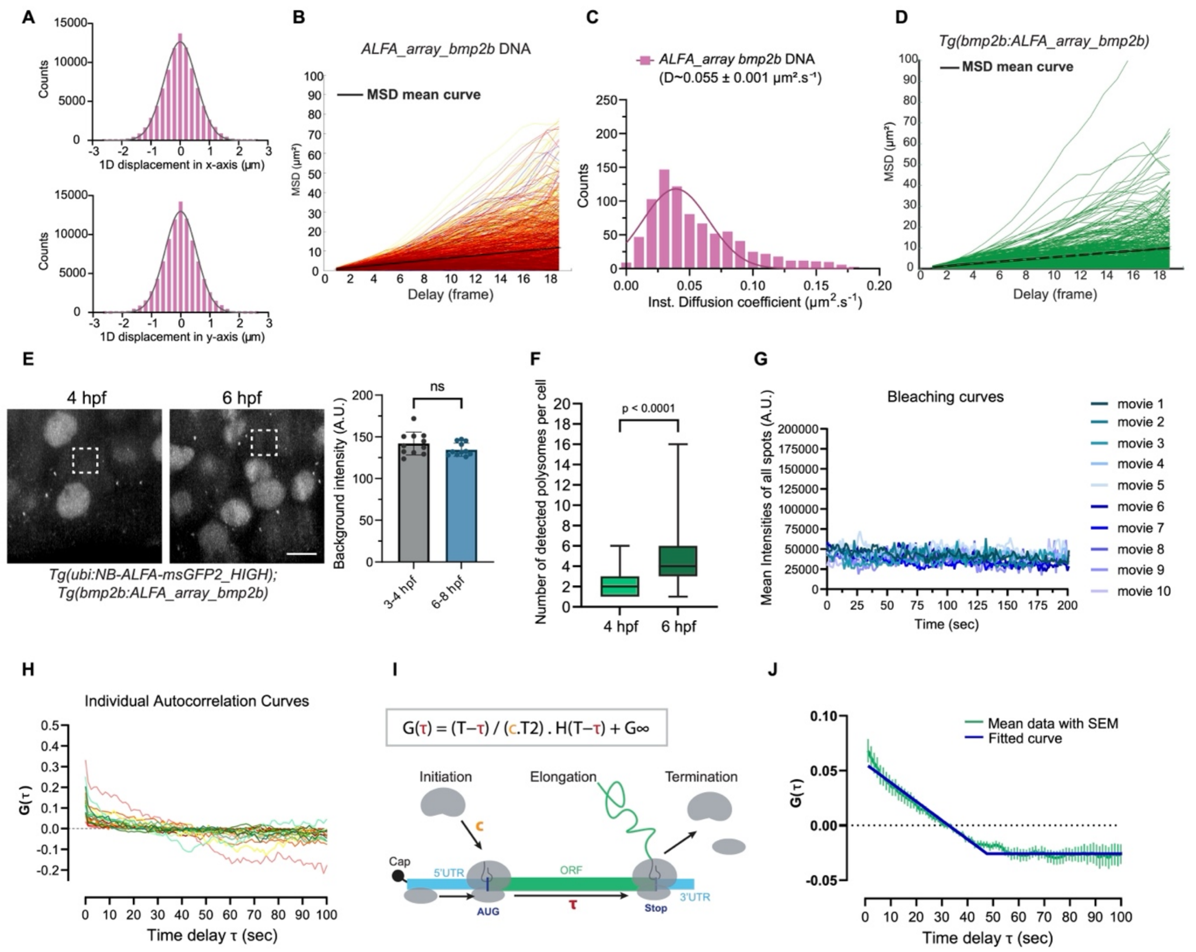


**fig. S5. Quantitative analysis of *bmp2b* polysome dynamics in zebrafish embryos.** (**A**) 1D displacement distribution in x and y of *bmp2b* polysomes. Gaussian fitting is shown with black curves. (**B**) Graph representing the Mean Square Displacement (MSD) curves as a function of time for translation particles of *bmp2b* (n=2729 traces, from 8 movies). Black curve represents the mean curve of all MSDs. (**C**) Frequency distribution of instantaneous diffusion coefficient of polysomes from *bmp2b* (n= 841 from 8 movies) measured from movies of 3-4.3 hpf *Tg(ubi:NB-ALFA-msGFP2_HIGH)* embryos injected with *ALFA_array_bmp2b* DNA. D is the estimated mean diffusion coefficient with the standard error of the mean (SEM). (**D**) Graph representing the Mean Square Displacement (MSD) curves as a function of time for *bmp2b* mRNA particles in translation from 3-4.3 hpf *Tg(ubi:NB-ALFA-msGFP2_HIGH); Tg(bmp2b:ALFA_array_bmp2b)* embryos (n=259 traces, from 12 movies). Black curve represents the mean curve of all MSDs. (**E**) Maximum intensity projected images from LLS movies of *Tg(ubi:NB-ALFA-msGFP2_HIGH); Tg(bmp2b:ALFA_array_bmp2b)* embryos at 4 and 6 hpf. Dashed squares mark a representative area used to quantify background intensity over time. Quantification of the background intensity is on the right. ns: non-significant with unpaired *t* test (p-value >0.05). (**F**) Quantification of the number of translation foci (polysomes) detected per cell from ~4 and ~6 hpf movies. (**G**) Representative control curve for bleaching correction of translation spots during time (seconds) from each movie of *Tg(ubi:NB-ALFA-msGFP2_HIGH); Tg(bmp2b:ALFA_array_bmp2b)* embryos. (**H**) Examples of autocorrelation curves $G(\tau)$ from single *bmp2b_AFLA_array* polysomes. (**I**) Equation used to fit the autocorrelation curves to retrieve the estimated parameters (see methods). ORF: Open Reading Frame. (**J**) Mean autocorrelation curve (green) from all movies (n=43 traces from 11 movies) with its Standard Error of Mean (SEM) and the fit (dark blue). Scale bar: 10 µm.

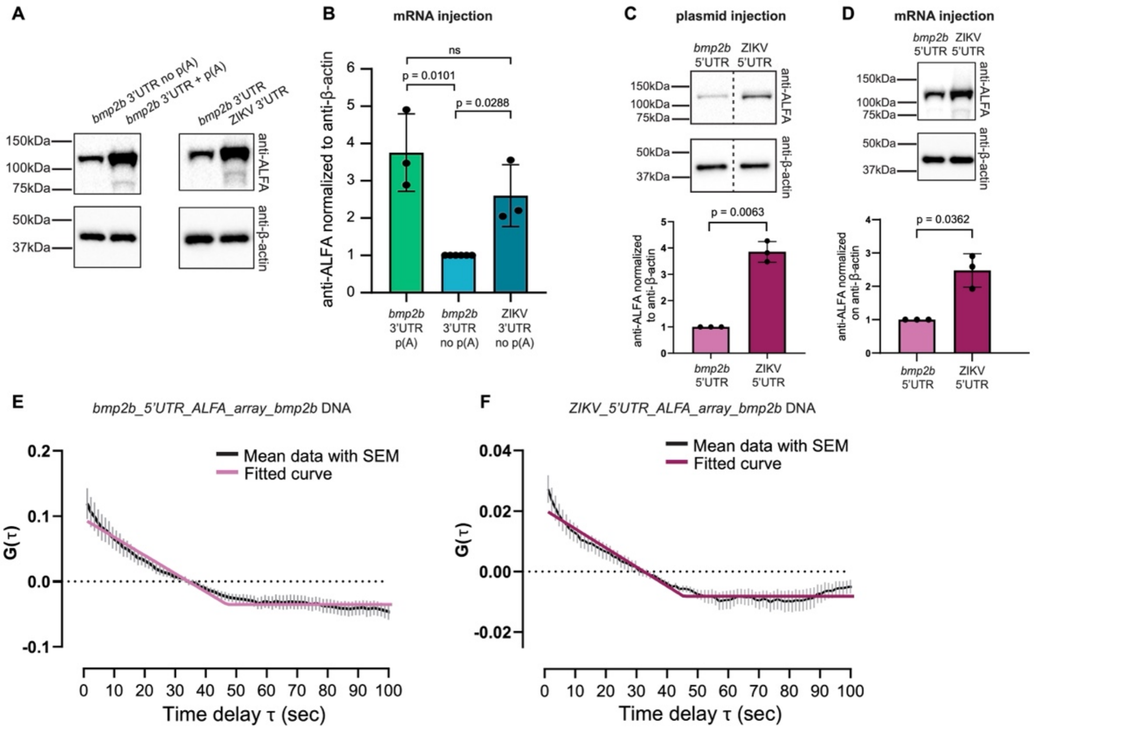


**fig. S6. Regulation of translation efficiency by *bmp2b* and ZIKV 5′ and 3′ untranslated regions.** (**A**) Western blot analysis, using anti-ALFA and anti-β-actin antibodies, of WT embryos injected with *bmp2b* 3'UTR *ALFA_array_bmp2b* mRNA with and without poly(A) tail; quantification from three independent crosses and injections. Two-tailed Welch's *t* test was performed. (**B**) Western blot analysis, using anti-ALFA and anti-β-actin antibodies, of WT embryos injected with *bmp2b* 3'UTR or with ZIKV 3'UTR *ALFA_array_bmp2b* mRNAs without poly(A) tail; quantification from three independent crosses and injections. Two-tailed Welch's *t* test was performed. (**C**) Western blot analysis, using anti-ALFA and anti-β-actin antibodies, of WT embryos injected with *bmp2b* 5'UTR or ZIKV 5'UTR *ALFA_array_bmp2b* DNA; quantification from three independent crosses and injections. Two-tailed Welch's *t* test was performed. (**D**) Western blot analysis, using anti-ALFA and anti-β-actin antibodies, of WT embryos injected with *bmp2b* 5'UTR or ZIKV 5'UTR *ALFA_array_bmp2b* mRNAs; quantification from three independent crosses and injections. Two-tailed Welch's *t* test was performed. (**E**) Mean autocorrelation curve (black) from *bmp2b* 5'UTR *ALFA_array_bmp2b* (n= 45 from 6 movies) mRNAs movies with its Standard Error of Mean (SEM) and the fit (pink) (see methods). (**F**) Mean autocorrelation curve (black) from ZIKV 5'UTR *ALFA_array_bmp2b* (n= 51 from 6 movies) mRNAs movies with its Standard Error of Mean (SEM) and the fit (dark pink) (see methods).

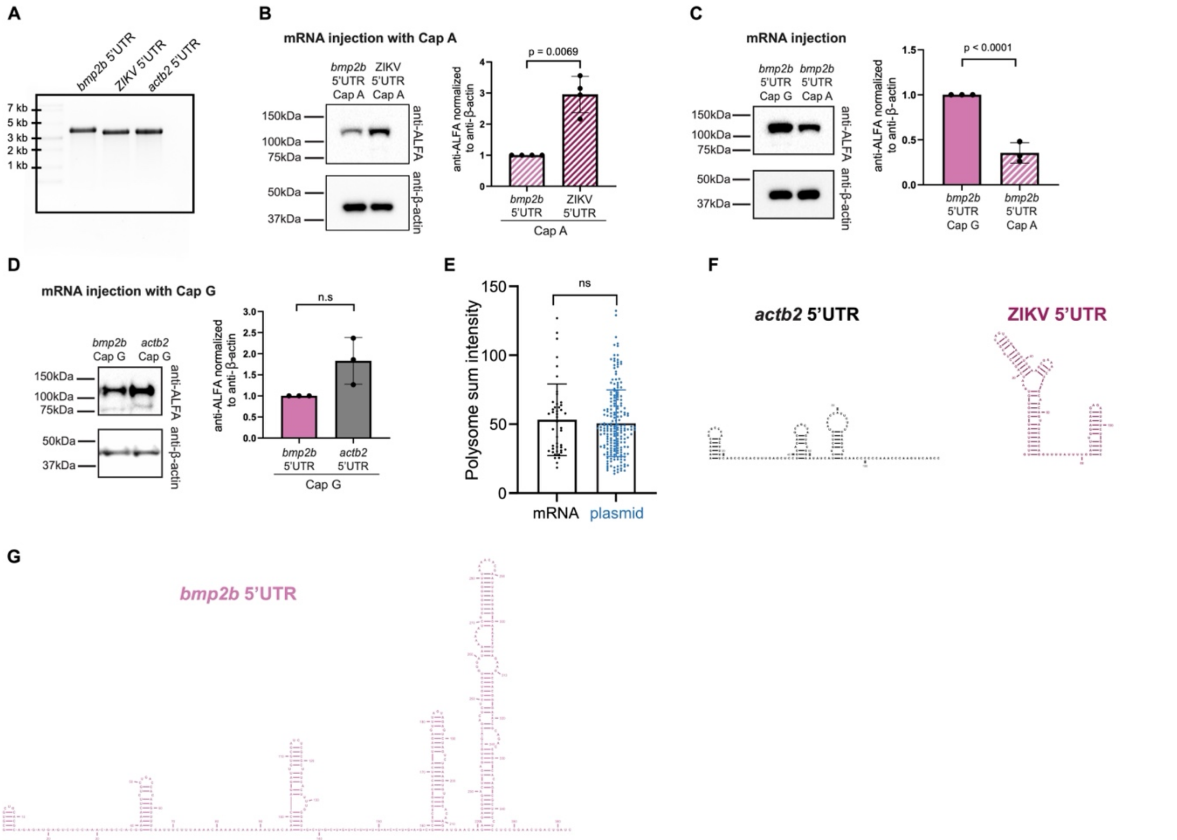


**fig. S7. Analysis of cap dependency in 5′UTR-driven translation of *ALFA_array_bmp2b* mRNA.** (**A**) Gel analysis confirming the expected size of in vitro-transcribed mRNAs containing the *bmp2b*, ZIKV, or *actb2* 5′UTR upstream of the *ALFA_array_bmp2b*. (**B**) Western blot analysis, using anti-ALFA and anti-β-actin antibodies, of WT embryos injected with *bmp2b* 5'UTR or ZIKV 5'UTR *ALFA_array_bmp2b* mRNAs with the cap analog A(ppp)G (Cap A); quantification from three 3 independent crosses and injections. Two-tailed Welch's *t* test was performed. (**C**) Western blot analysis, using anti-ALFA and anti-β-actin antibodies, of WT embryos injected with *ALFA_array_bmp2b* mRNAs with $m^7G$(ppp)G (Cap G) or A(ppp)G (Cap A) cap; quantification from 3 independent crosses and injections. Two-tailed Welch's *t* test was performed. (**D**) Western blot analysis, using anti-ALFA and anti-β-actin antibodies, of WT embryos injected with *bmp2b* 5'UTR or *actb2* 5'UTR *ALFA_array_bmp2b* mRNAs with $m^7G$(ppp)G (Cap G) cap; quantification from three independent crosses and injections. ns: non-significant with a two-tailed Welch's *t* test (p-value >0.05). (**E**) Polysome sum intensities (normalized to background) of *ALFA_array_bmp2b* mRNAs derived from in vitro-transcribed mRNAs (n = 45 from 3 movies) or plasmid DNA (n = 200 from 4 movies). ns: non-significant with a two-tailed Welch's *t* test. (**F**) Predicted secondary structure of the *actb2* and ZIKV 5'UTR. (**G**) Predicted secondary structure of the *bmp2b* 5'UTR.

**Table S1.**

| Plasmid: | Sequence: |
|---|---|
| **NB-msGFP2**<br><br>Ubiquitin promoter > NB-ALFA - msGFP2 - GB1NLS - SV40 | accagcaaagttctagaatttgtcgaaacatttatgttatatatttcctgaaaaaaattctgagtaagttcttaagtgttattgccagcaacataaacaacagacggcaaaatgaataaatgataaca<br>aagcagtaggcttaaataaacctaatttttataggctgttctctacaaccctcaaacagtgattagttttgtacttataaacttgccctttcattcatatttcaagaaaattggttcagaagatctggata<br>ttctagcagttgttcaagctcatggagggatcagtgacctgattccacaatgactaggcctaatccagaaattagatgactgtcaacataaaaaggcacagcactcactagctgccctatatattt<br>tattatattttacatatattattttatttatttagctctgagtgctgtactttctggttaaagaaaactgcttacaacagctaacctgtactacctcaggctcagggaatttggaacaggtttgtctggtttg<br>tttctttaaccatgcatgcttgttttcaactatggcaacacagtcacatgggacattacagaaatgatttgtcgatgacatgcgacttttctttaataaagcgcaaagatcccaaaaagcaaactttta<br>acaaaaatcatataattatattttcaatccagctttgtagcaactttgtgctgctgttcactcagcaacagatagtcagtataaggtcagtgtgtctcaaagcagtgccatctgtttcacacattgcgtt<br>ctatatataagtgtgctggttgacacgacactgtataaggcctaggctaaaacacaaacaatgtagaatgacactgtgttttttttgtaaacaaatgttgtttttggttaaacatctttgtgaaaacatc<br>ctcctgtcatgtatttgctatattcaaatgttaaacccgtgcagaatagaacatatacaaaaaaaaacaacacaacacatttttaaacattattaaatatcaagtattgctggcagttctgtttctgtttt<br>acagtaccctttgccacagttctccgcttttcctggtccagattccacaagtctgattcaccaatagcaaagcgaataaacaaccaaagcagccaatcactgcttgtagactgtcctgcgagac<br>cggcccattccagcacattctggaaacttcctttatatgataattataaatacatttaaattattgatacaaaacatgtaattcctagaacataaccatagcaatcattagttttcagggtaattatgtatt<br>tttaggatttgactgcggaaagatctggtcatgtgacgtctcatgaacgtcacggccctgggtttctataaatacagtaggactctcgaccatcggcagatttttcgaagaagaagatcagtttca<br>ggagccgtactgttccgtttcaacgcaaatattaacggtaagagcgaatttcctagtttgttttcatgccattctttaaaaccatagcgtattactttaattatagtaaactttcgctttctttattacaag<br>agacgttttgtgttgattctccgcggacattttcggtcagacaatcagaaaatgaccgcggaggaccagtaacttgcattacacgtaagttaaatcttcgtgtattaaaatggttaggttgttaacg<br>tcaaataggttaccgtgtttgcgtgtgatcaggttggttttgttagatttttgtcagtatttttaatttatttgttttagtttatttatttttttgctgaatcatagtttgtgaacaaagaacccggatgttacat<br>acagtacagccgccatgttacagagagttataacttaatcattttaaaaataattttgccttacttttagtttgtcatgttgagaaatgaggaaatgttaaaatgaggaaatatccaattaatttaatata<br>tcaaaataatccatgattacaatgcactgaactggagaaaattaagatgttttctagtgtcatgaaacaaatgtaagagatgtacattgtagatgttttatgtcaagaattggctagttgatgcagca<br>tactggcgatactcagttgtaataacagtaacgttacatgttaatagactactgagtatgctgttctgtctatgtatgctctgtaagctacgagaaggactttttaaacagtaaagggtgcaatatttt<br>tacaaattgaattaaataaaggctgtctattaagtaatatgcttgatatttttcttacttgatcggaaataagaaaaaatataaacgttgttgctctaaaaatcctagttcagtttagccaaccacaaata<br>ccttttgttcctccaacagttttttttttcttctctataatatttggcagtctatagtactccaaatgtttccccacagtctaactaattggtacagccaaaatcatgacacttattgcaataataattttggtt<br>cattggcattgttgatagcctgtgccactaatatggtcgattgatcatgcttcaggaagaaaactatattgtttgatgtaagattattaaatcttcacctgcttccattacaaactattcccatcttattga<br>attctggtatgtcttaaaggattagttcacttcccaaatcaaaatttacttagcctttttcatccatgatcccttttttttcatcattaatgaagaaattgtttttgaaaaagtttcaagatttttttctctatattg<br>tggagcttgttagtttaaaattccaaaatgcaatatgtggcttcaaatggttctaaatgatcccagtcaaggaataacagtcttatctaatgaaaccattagacctttaaaaataaaaaaaaataaaag<br>tatttatttttaaatgactgagtgattaagttgaatttcagcgtttccttactgtgtagaagtccttccttactggccccaccctttggttctctgccaatctgctacctaatgtaatgttgtggaacattatt<br>attctttatttcttaatttttttatttttttattttaaaacaatgtaaactgcacagatgtgcagtttgtttaaaatggccaatgctttggaaatgcatgacataattagatttcatgatgcacaaagccaaatct<br>cagagcttgtgcaaaatgagctatcatttcactaggtaagaccctaaattttcatataggatcatttggacaattttgctgcaggtaaaatgcattctatagtccactgtcagccattgttttggatagt<br>atttatttttctctacaagtatagtcaatagttttctattattttaaaggtttgtaacatttaagggtgaccaaatgcaaagtaaaatttcattttcgggtgaactatctcgtttaacatgggagaagtgca<br>aaacatacattattggctagaacattgtagtatttttttaaatggaaatgtgtgattgctaatcttactttgaatttgtttacagggccggccatggaggtgcagctgcaggagagcggaggaggtct<br>ggtgcaaccaggaggcagcttgcgtttgagctgtacggccagcggcgtcaccatcagtgccctcaatgcaatggccatgggctggtatcgccaggcgcccggcgaaagacgagtgatg<br>gtcgctgctgtatcggagcgtggcaatgccatgtatcgggaatcggttcaaggacgctttacggttacaagggatttcaccaacaagatggtgtcccttcagatggacaacctaaaaccgga<br>ggacacagcggtctactactgccatgtactggaggatcgcgtggattccttccacgactactggggccagggcacccaggtgactgtttcctctggtggtggcggcagcggcggtggcgg<br>cagtggtgggggcggttcaggcggaggaggatcggatagcacggagagtctgttcaccggggtggtgcccatcctggtcgagctggacggcgacgtaaacggccacaagttcagcgtg<br>cgcggcgagggcgagggcgatgccaccaacggcaagctgaccctgaagttcatctgcaccaccggcaagctgcccgtgccctggcccaccctcgtgaccaccctgacctacggcgtg<br>cagtgcttcagccgctaccccgaccacatgaagcagcacgacttcttcaagtccgccatgcccgaaggctacgtccaggagcgcaccatcaccttcaaggacgacggcacctacaagac<br>ccgcgccgaggtgaagttcgagggcgacaccctggtgaaccgcatcgagctgaagggcatcgacttcaaggaggacggcaacatcctggggcacaagctggagtacaactacaacag<br>ccacaacgtctatatcacggccgacaagcagaagaacggcatcaaggcgaacttcaagatccgccacaacgtcgaggacggcagcgtgcagctcgccgaccactaccagcagaacac<br>ccccatcggcgacggccccgtgctgctgcccgacaaccactacctgagcacccagtccaagctgagcaaagaccccaacgagaagcgcgatcacatggtcctgctggagttcgtgacc<br>gccgccgggatcactggaggctcaggaggaagcggtggaggtcggaccgaagagtacaagcttatcctgaacggtaaaaccctgaaaggtgaaaccaccaccgaagctgttgacgctg<br>ctaccgcggaaaaagttttcaaacagtacgctaacgacaacggtgttgacggtgaatggacctacgacgacgctaccaaaaccttcacggtaaccgaaggtggtggtagcggtggtggta<br>ctagtcccaagaagaagcgcaaggtgtaaggtaccatcgatgatgatccagacatgataagatacattgatgagtttggacaaaccacaactagaatgcagtgaaaaaaatgctttatttgtga<br>aatttgtgatgctattgctttatttgtaaccattataagctgcaataaacaagttaacaacaacaattgcattcattttatgtttcaggttcagggggaggtgtgggaggttttt |
| **scFv-msGFP2**<br><br>Ubiquitin promoter > scFv - msGFP2 - GB1NLS - SV40 | accagcaaagttctagaatttgtcgaaacatttatgttatatatttcctgaaaaaaattctgagtaagttcttaagtgttattgccagcaacataaacaacagacggcaaaatgaataaatgataaca<br>aagcagtaggcttaaataaacctaatttttataggctgttctctacaaccctcaaacagtgattagttttgtacttataaacttgccctttcattcatatttcaagaaaattggttcagaagatctggata<br>ttctagcagttgttcaagctcatggagggatcagtgacctgattccacaatgactaggcctaatccagaaattagatgactgtcaacataaaaaggcacagcactcactagctgccctatatattt<br>tattatattttacatatattattttatttatttagctctgagtgctgtactttctggttaaagaaaactgcttacaacagctaacctgtactacctcaggctcagggaatttggaacaggtttgtctggtttg<br>tttctttaaccatgcatgcttgttttcaactatggcaacacagtcacatgggacattacagaaatgatttgtcgatgacatgcgacttttctttaataaagcgcaaagatcccaaaaagcaaactttta<br>acaaaaatcatataattatattttcaatccagctttgtagcaactttgtgctgctgttcactcagcaacagatagtcagtataaggtcagtgtgtctcaaagcagtgccatctgtttcacacattgcgtt<br>ctatatataagtgtgctggttgacacgacactgtataaggcctaggctaaaacacaaacaatgtagaatgacactgtgttttttttgtaaacaaatgttgtttttggttaaacatctttgtgaaaacatc<br>ctcctgtcatgtatttgctatattcaaatgttaaacccgtgcagaatagaacatatacaaaaaaaaacaacacaacacatttttaaacattattaaatatcaagtattgctggcagttctgtttctgtttt<br>acagtaccctttgccacagttctccgcttttcctggtccagattccacaagtctgattcaccaatagcaaagcgaataaacaaccaaagcagccaatcactgcttgtagactgtcctgcgagac<br>cggcccattccagcacattctggaaacttcctttatatgataattataaatacatttaaattattgatacaaaacatgtaattcctagaacataaccatagcaatcattagttttcagggtaattatgtatt<br>tttaggatttgactgcggaaagatctggtcatgtgacgtctcatgaacgtcacggccctgggtttctataaatacagtaggactctcgaccatcggcagatttttcgaagaagaagatcagtttca<br>ggagccgtactgttccgtttcaacgcaaatattaacggtaagagcgaatttcctagtttgttttcatgccattctttaaaaccatagcgtattactttaattatagtaaactttcgctttctttattacaag<br>agacgttttgtgttgattctccgcggacattttcggtcagacaatcagaaaatgaccgcggaggaccagtaacttgcattacacgtaagttaaatcttcgtgtattaaaatggttaggttgttaacg<br>tcaaataggttaccgtgtttgcgtgtgatcaggttggttttgttagatttttgtcagtatttttaatttatttgttttagtttatttatttttttgctgaatcatagtttgtgaacaaagaacccggatgttacat<br>acagtacagccgccatgttacagagagttataacttaatcattttaaaaataattttgccttacttttagtttgtcatgttgagaaatgaggaaatgttaaaatgaggaaatatccaattaatttaatata<br>tcaaaataatccatgattacaatgcactgaactggagaaaattaagatgttttctagtgtcatgaaacaaatgtaagagatgtacattgtagatgttttatgtcaagaattggctagttgatgcagca<br>tactggcgatactcagttgtaataacagtaacgttacatgttaatagactactgagtatgctgttctgtctatgtatgctctgtaagctacgagaaggactttttaaacagtaaagggtgcaatatttt<br>tacaaattgaattaaataaaggctgtctattaagtaatatgcttgatatttttcttacttgattggaaataagaaaaaaatataaacgttgttgctctaaaaatcctagttcagtttagccaaccacaaata<br>ccttttgttcctccaacagttttttttttcttctctataatatttggcagtctatagtactccaaatgtttccccacagtctaactaattggtacagccaaaatcatgacacttattgcaataataattttggtt<br>cattggcattgttgatagcctgtgccactaatatggtcgattgatcatgcttcaggaagaaaactatattgtttgatgtaagattattaaatcttcacctgcttccattacaaactattcccatcttattga<br>attctggtatgtcttaaaggattagttcacttcccaaatcaaaatttacttagcctttttcatccatgatcccttttttttcatcattaatgaagaaattgtttttgaaaaagtttcaagatttttttctctatattg<br>tggagcttgttagtttaaaattccaaaatgcaatatgtggcttcaaatggttctaaatgatcccagtcaaggaataacagtcttatctaatgaaaccattagacctttaaaaataaaaaaaaataaaag<br>tatttatttttaaatgactgagtgattaagttgaatttcagcgtttccttactgtgtagaagtccttccttactggccccaccctttggttctctgccaatctgctacctaatgtaatgttgtggaacattatt<br>attctttatttcttaatttttttatttttttattttaaaacaatgtaaactgcacagatgtgcagtttgtttaaaatggccaatgctttggaaatgcatgacataattagatttcatgatgcacaaagccaaatct<br>cagagcttgtgcaaaatgagctatcatttcactaggtaagaccctaaattttcatataggatcatttggacaattttgctgcaggtaaaatgcattctatagtccactgtcagccattgttttggatagt<br>atttatttttctctacaagtatagtcaatagttttctattattttaaaggtttgtaacatttaagggtgaccaaatgcaaagtaaaatttcattttcgggtgaactatctcgtttaacatgggagaagtgca<br>aaacatacattattggctagaacattgtagtatttttttaaatggaaatgtgtgattgctaatcttactttgaatttgtttacagggccggatgggccccgacatcgtgatgacccagagccccagca<br>gcctgagcgccagcgtgggcgaccgcgtgaccatcacctgccgcagcagcaccggcgccgtgaccaccagcaactacgccagctgggtgcaggagaagcccggcaagctgttcaag<br>ggcctgatcggcggcaccaacaaccgcgcccccggcgtgcccagccgcttcagcggcagcctgatcggcgacaaggccaccctgaccatcagcagcctgcagcccgaggacttcgc<br>cacctacttctgcgccctgtggtacagcaaccactgggtgttcggccagggcaccaaggtggagctgaagcgcggcggcggcggcagcggcggcggcggcagcggcggcggcggc<br>agcagcggcggcggcagcgaggtgaagctgctggagagcggcggcggcctggtgcagcccggcggcagcctgaagctgagctgcgccgtgagcggcttcagcctgaccgactacg |

| | |
|---|---|
| | gcgtgaactgggtgcgccaggcccccggccgcggcctggagtggatcggcgtgatctggggcgacggcatcaccgactacaacagcgccctgaaggaccgcttcatcatcagcaagg<br>acaacggcaagaacaccgtgtacctgcagatgagcaaggtgcgcagcgacgacaccgccctgtactactgcgtgaccggcctgttcgactactggggccagggcaccctggtgaccgt<br>gagcagctacccatacgatgttccagattacgctggtggaggcggaggttctgggggaggaggtagtggcggtggtggttcaggaggcggcggaagcttggatccaggtggaggtgga<br>agcggtgatagcacggagagtctgttcaccggggtggtgcccatcctggtcgagctggacggcgacgtaaacggccacaagttcagcgtgcgcggcgagggcgagggcgatgccacc<br>aacggcaagctgaccctgaagttcatctgcaccaccggcaagctgcccgtgccctggcccaccctcgtgaccaccctgacctacggcgtgcagtgcttcagccgctaccccgaccacatg<br>aagcagcacgacttcttcaagtccgccatgcccgaaggctacgtccaggagcgcaccatcaccttcaaggacgacggcacctacaagacccgcgccgaggtgaagttcgagggcgaca<br>ccctggtgaaccgcatcgagctgaagggcatcgacttcaaggaggacggcaacatcctggggcacaagctggagtacaactacaacagccacaacgtctatatcacggccgacaagca<br>gaagaacggcatcaaggcgaacttcaagatccgccacaacgtcgaggacggcagcgtgcagctcgccgaccactaccagcagaacacccccatcggcgacggccccgtgctgctgcc<br>cgacaaccactacctgagcacccagtccaagctgagcaaagaccccaacgagaagcgcgatcacatggtcctgctggagttcgtgaccgccgccgggatcactggaggctcaggagga<br>agcggtggaggtcggaccgaagagtacaagcttatcctgaacggtaaaaccctgaaaggtgaaaccaccaccgaagctgttgacgctgctaccgcggaaaaagttttcaaacagtacgct<br>aacgacaacggtgttgacggtgaatggacctacgacgacgctaccaaaaccttcacggtaaccgaaggtggtggtagcggtggtggtactagtcccaagaagaagcgcaaggtgtaagt<br>accatcgatgatgatccagacatgataagatacattgatgagtttggacaaaccacaactagaatgcagtgaaaaaaatgctttatttgtgaaatttgtgatgctattgctttatttgtaaccattata<br>agctgcaataaacaagttaacaacaacaattgcattcattttatgtttcaggttcagggggaggtgtgggaggtttttt |
| **NB-mScarlet3**<br><br>Ubiquitin promoter > NB-ALFA – mScarlet3 - GB1 - SV40 | accagcaaagttctagaatttgtcgaaacatttatgttatatatttcctgaaaaaaattctgagtaagttcttaagtgttattgccagcaacataaacaacagacggcaaaatgaataaatgataaca<br>aagcagtaggcttaaataaacctaatttttataggctgttctctacaaccctcaaacagtgattagttttgtacttataaacttgccctttcattcatatttcaagaaaattggttcagaagatctggata<br>ttctagcagttgttcaagctcatggagggatcagtgacctgattccacaatgactaggcctaatccagaaattagatgactgtcaacataaaaaggcacagcactcactagctgccctatatattt<br>tattatattttacatatattattttatttatttagctctgagtgctgtactttctggttaaagaaaactgcttacaacagctaacctgtactacctcaggctcagggaatttggaacaggtttgtctggtttg<br>tttctttaaccatgcatgcttgttttcaactatggcaacacagtcacatgggacattacagaaatgatttgtcgatgacatgcgacttttctttaataaagcgcaaagatcccaaaaagcaaactttta<br>acaaaaatcatataattatattttcaatccagctttgtagcaactttgtgctgctgttcactcagcaacagatagtcagtataaggtcagtgtgtctcaaagcagtgccatctgtttcacacattgcgtt<br>ctatatataagtgtgctggttgacacgacactgtataaggcctaggctaaaacacaaacaatgtagaatgacactgtgttttttttgtaaacaaatgttgtttttggttaaacatctttgtgaaaacatc<br>ctcctgtcatgtatttgctatattcaaatgttaaacccgtgcagaatagaacatatacaaaaaaaaacaacacaacacatttttaaacattattaaatatcaagtattgctggcagttctgtttctgtttt<br>acagtaccctttgccacagttctccgcttttcctggtccagattccacaagtctgattcaccaatagcaaagcgaataaacaaccaaagcagccaatcactgcttgtagactgtcctgcgagac<br>cggcccattccagcacattctggaaacttcctttatatgataattataaatacatttaaattattgatacaaaacatgtaattcctagaacataaccatagcaatcattagttttcagggtaattatgtatt<br>tttaggatttgactgcggaaagatctggtcatgtgacgtctcatgaacgtcacggccctgggtttctataaatacagtaggactctcgaccatcggcagatttttcgaagaagaagatcagtttca<br>ggagccgtactgttccgtttcaacgcaaatattaacggtaagagcgaatttcctagtttgttttcatgccattctttaaaaccatagcgtattactttaattatagtaaactttcgctttctttattacaag<br>agacgttttgtgttgattctccgcggacattttcggtcagacaatcagaaaatgaccgcggaggaccagtaacttgcattacacgtaagttaaatcttcgtgtattaaaatggttaggttgttaacg<br>tcaaataggttaccgtgtttgcgtgtgatcaggttggttttgttagatttttgtcagtatttttaatttatttgttttagtttatttattttttttgctgaatcatagtttgtgaacaaagaacccggatgttacat<br>acagtacagccgccatgttacagagagttataacttaatcattttaaaaataattttgccttacttttagtttgtcatgttgagaaatgaggaaatgttaaaatgaggaaatatccaattaatttaatata<br>tcaaaataatccatgattacaatgcactgaactggagaaaattaagatgttttctagtgtcatgaaacaaatgtaagagatgtacattgtagatgttttatgtcaagaattggctagttgatgcagca<br>tactggcgatactcagttgtaataacagtaacgttacatgttaatagactactgagtatgctgttctgtctatgtatgctctgtaagctacgagaaggactttttaaacagtaaagggtgcaatatttt<br>tacaaattgaattaaataaaggctgtctattaagtaatatgcttgatatttttcttacttgatcggaaataagaaaaaatataaacgttgttgctctaaaaatcctagttcagtttagccaaccacaaata<br>ccttttgttcctccaacagttttttttcttctctataatatttggcagtctatagtactccaaatgtttccccacagtctaactaattggtacagccaaaatcatgacacttattgcaataataattttggtt<br>cattggcattgttgatagcctgtgccactaatatggtcgattgatcatgcttcaggaagaaaactatattgtttgatgtaagattattaaatcttcacctgcttccattacaaactattcccatcttattga<br>attctggtatgtcttaaaggattagttcacttcccaaatcaaaatttacttagcctttttttcatccatgatccctttttttcatcattaatgaagaaattgtttttgaaaaagtttcaagatttttttctctatattg<br>tggagcttgttagtttaaaattccaaaatgcaatatgtggcttcaaatggttctaaatgatcccagtcaaggaataacagtcttatctaatgaaaccattagacctttaaaaataaaaaaaataaaag<br>tatttatttttaaatgactgagtgattaagttgaatttcagcgtttccttactgtgtagaagtccttccttactggccccaccctttggttctctgccaatctgctacctaatgtaatgttgtggaacattatt<br>attctttatttcttaatttttttatttttttattttaaaacaatgtaaactgcacagatgtgcagtttgtttaaaatggccaatgctttggaaatgcatgacataattagatttcatgatgcacaaagccaaatct<br>cagagcttgtgcaaaatgagctatcatttcactaggtaagaccctaaattttcatataggatcatttggacaattttgctgcaggtaaaatgcattctatagtccactgtcagccattgttttggatagt<br>atttatttttctctacaagtatagtcaatagttttctattattttaaaggtttgtaacatttaagggtgaccaaatgcaaagtaaaatttcattttcgggtgaactatctcgtttaacatgggagaagtgca<br>aaacatacattattggctagaacattgtagtatttttttaaatggaaatgtgtgattgctaatcttactttgaatttgtttacagggccggatggaggtgcagctgcaggagagcggaggaggtctgg<br>tgcaaccaggaggcagcttgcgtttgagctgtacggccagcggcgtcaccatcagtgccctcaatgcaatggccatgggctggtatcgccaggcgcccggcgaaagacgagtgatggtc<br>gctgctgtatcggagcgtggcaatgccatgtatcgggaatcggttcaaggacgctttacggttacaagggatttcaccaacaagatggtgtcccttcagatggacaacctaaaaccggagga<br>cacagcggtctactactgccatgtactggaggatcgcgtggattccttccacgactactggggccagggcacccaggtgactgtttcctctggtggtggcggcagcggcggtggcggcagt<br>ggtgggggcggttcaggcggaggaggatcggatagcaccgaggcagtgatcaaggagttcatgcggttcaaggtgcacatggagggctccatgaacggccacgagttcgagatcgag<br>ggcgagggcgagggccgcccctacgagggcacccagaccgccaagctgagggtgaccaagggtggccccctgcccttctcctgggacatcctgtcccctcagttcatgtacggctcca<br>gggccttcacgaagcaccccgccgacatccccgactactggaagcagtccttccccgagggcttcaagtgggagcgcgtgatgaacttcgaggacggcggcgccgtgtccgtggccca<br>ggacacctccctggaggacggcaccctgatctacaaggtgaagctccgcggcaccaacttccctcctgacggccccgtaatgcagaagaagacaatgggctgggaagcatccaccgag<br>cggttgtaccccgaggacgtcgtgctgaagggcgacattaagatggccctgcgcctgaaggacggcggccgctacctggcggacttcaagaccacctacagggccaagaagcccgtgc<br>agatgcccggcgccttcaacatcgaccgcaagttggacatcacatcccacaacgaggactacaccgtggtggaacagtacgaacgctccgtggcccgccactccaccggcggctccggt<br>ggctccggtggaggtcggaccgaagagtacaagcttatcctgaacggtaaaaccctgaaaggtgaaaccaccaccgaagctgttgacgctgctaccgcggaaaaagttttcaaacagtac<br>gctaacgacaacggtgttgacggtgaatggacctacgacgacgctaccaaaaccttcacggtaaccgaaggtggtggtagcggtggtggtactagttaagtaccatcgatgatgatccaga<br>catgataagatacattgatgagtttggacaaaccacaactagaatgcagtgaaaaaaatgctttatttgtgaaatttgtgatgctattgctttatttgtaaccattataagctgcaataaacaagttaa<br>caacaacaattgcattcattttatgtttcaggttcagggggaggtgtgggaggtttttt |
| ***bmp2b_ALFA_array***<br><br>bmp2b promoter (5'UTR) > ALFA_array - bmp2b (intron) - 3'UTR | cattgctcctcttcccagtttggtcgagggttaacataatatcaagtcgtgcatttcctattgacaatatagcatgtttacctgtggattgatgtataaaacaagttcgactgatttatgcaaagcaaa<br>cacactgttgtgttacagcaaccttctgacacatcagcaatcgatacacacagagcattgttatttaaatcagacttctccagaacagtgtctgctcaggatttaacttgatagtgtgaactgattgt<br>gatgcaagatgtgttggaaatatatatataaagtaatgtatacttttacactattttacagcacaattaaacagattttctattagcacaggggttttaaaactcccaagggccgtaaaaggggtgct<br>gtcttgtgtttttctatagagggaggctactgttgcctgtaggctatgtcttcaggaaaagggtagtttgaacattgaagtaaatattactatgaatttcattaggtgaaaatgtttcatgtgtaacattt<br>atacaagcaactgcactttaatattgcgggaaaatgtaaatgtgtttactattttaatactaaacatatgcaattataattgactctataaagactctataaattgttgtgagcatcacattttggacaaa<br>tcaaaagtatccggaaaatataatgcatacctgtacactttgagttcttttttttttttttgaacacaggagctttaaaactgagggccggggactcccaggagtccataaagtggtgccatcttttc<br>ttttcttcttttttcttctgcttgtctagccagtggcaataatctccctcaggcaatattttctgtaaagctgctaaagaaaaaagtggcatagaaataaatgttacatatatatatatacatacacagtg<br>ttcatcatatataagtacaccctttgaaaatctctcaattcatattttgtgcatatactttagataagtcagtactgaagccaaatctgaagctaatctaagaaaataacatgatagtctaagttagtac<br>actcaaatttatgtaagggaaaatattaagtaaaaaatgtaaaaagagcacaattcatatatgtatattgtttatatatgtatacatataggaagctttacaatattatatttgtgcatatacattagacta<br>gtcagtattaaagccaaatctggagctaatttaacaaaataacttatgattatagtttaaaatttgtacacgcaaatttataagttaagcaaatatatatatatatatatatatatatatacatacatac<br>atacatacatacctcaattcaaggtatttttatttatttatttatttttgttaactaactgtaaacagaattattagccccccttgtttatttttttcccccatttcttctgtttaacggaaagcagattttttttaagc<br>agactttaaaatgtttctttaaaaataaaaaacttgttattttctagccaaaataaaacaaataagactttctcccttgctctgataaaaatcatttgggaaatataaaaaaaaagaacaaaatttcaaa<br>ggggcactaataattctgacatcaactttaaattttatttatttatcttttggtaactacgacaaaaagagatgtaatttagctttatagctaactctggcataacatgtcatgttgtagctacattgtccc<br>agaataagtaaataaatgaatattcggctttatacaagtctaaaatagttgtacataaaatattagatcattttaaaacgtttaaagacaacacattgcaataacaaatcaattaaatgaaacctaaa<br>ataacgttaacatttttccttcactataaattactatacatgattttaaacagaagagatatccttataaatactgaaaaaaatactcaaatacaaatgtagataatttaaattagtgcgcatttaaattta<br>ggatttgtttaaccatacttcagtctcaattgtattgcgtatacattacgttctcgttcaaattactaacatgtttacataggataatgcataaaatatgccccatgcaggggaaattcggtccatccgc<br>gcgcgcagagtgtgggcatgttcaaacgcttgaatggagagagcgcggcatcattgtgacatcatcagacaacaaaaagcctcgcgctcgcgcagcgaagcgctccaatcaatggcaca<br>gacgcggcgcgtgctgcacgcagagatgagtctccaaacagccacggaaaacttctgctgaccacaagttttcgatttctttaaaacaaaaacaaaaaatgacaaatccaggattgtgcgat<br>ctcgcgctgtcacttttgggattgctgctgtctttgacctgagcgctcgcgcacttcattagagtttagtagagtctagtctgaagtgttgcacaagtatgaacaagaagaggcgacttgagctgc<br>gacgactctctgtcgtgggataaaaaaatcgcttgtggattaaaacacgaattcatgaggaacttagaagacgacgggaacgcagaccggccacagcgcttcctcctccggtaacgcattc<br>aattacatgcttgatattacgtattagcaaacagtgcgactgttttaaaaactaatgctgacatttctgtaactttgattgcacttcacatcactgcgtgtgcgtgcctgcatgtaatgtgtgaggtca<br>ggaggtttgcgcaaatgaaaagaaaacgcttctgcgcttccttatcatcagatctctttgtcgtctttgttgcccgagttgtttttgcgtaagatcgaccacctgtagcctcacaactctgcctctcc |

| | |
|---|---|
| | cttagcgaaatgtcttttgctagacattcagtccttttgaagactttcctttcatgatctaaagacagacccccacccagacagcgcgctcttatctccctttaattgcagactccatgtctgctgtctt<br>catgtctcccttctttcgatttcttttgattcactctttcctgtttcatcccatttgtctgacatcctctttggagtctcccgctttcactgtagatcaatctcttattatagcctgaggcagggatattattctt<br>aatctttttttagagcgcgagtgtgtgttcatcgcctcaaagacatctccaaaagatccagatatgtctgtggtttgaggcagccgcaggttgaacgcggccagggaacttttccatgcgtataa<br>gtggttgaagctctgggcttgaagtcaaacgtgtttttttaggacttaaaggcgcaggtattccataaaatcaaactattatgagttattggcggattgaacctatctttaggtgatctatgagccatt<br>gtgccactgagcaaggcacttattaacgacaccagattgctccaagaggggctgtagtttaaaaatgtactcttcaaaaataaagttattttaaataattaaactttacttacagaatatattttcttg<br>caatatttaaacattcttaaatcaagaacatcacagttttgtttttagaaataagaagtgaaattaagtgaggctttctttaaaacaagctaaattatttccagtggggtaagtaaataatcttatgtcaa<br>actaaaagcaagaattgtttactggcttgatttgaggaaaaacacttgatttcgacgttttatttctgaaaacaaagcatttcacttgtttagaaaatgcaattcgatagatagatatttgatttttagat<br>atttacactagaaaccagacaaaaaccattttaattttcaggaaatgcatgtttttatgcagcgctttacattatttttactgttacgttacaggtttagatcaatgtaaagttattatagttattggtggat<br>tggacctatcttttagatgatctatgagccattgtgccactaagcaaggcacttaattgctccagagattgctccagaagggctgtagatttaagaaatgtactcttaaaaataaacttattttatgaa<br>ttaatcttcacgaaaaatatatattcttactctcgagtctaaatatttaaacattgtaaaatcaagaagcattttctaaacaagtaaggtccaaattaagtgaggatttctttaaagcaagctcaattatc<br>ttccagtagggtaagtaaaataatcttatttcaaactaaaagcaagattattttactcaccaaactggttagatttgaggaaaaacacataatttttacattttatttctgaaagcaaaacattgcacttg<br>tctatgaataatatgatgtaaatataatttaataaattattaataatttaaattatgaaattgaataataatacaataataataaaatattaatatatatatatatttacacctgaaacgagacaaaaactctt<br>tttggaaaaccacactttatgcagtgctatcatgtattttaattaaaatgttatattacaagttcagatcgatgtgttgctttggattctgtctcccttaaagcagcagagcatgcttttcccaagaaacc<br>agaaaataaaaagttcactgtgggccctaaaaaggttcatctctatgatgcaatgctgtaaaacctttaaaatgatttatggtaagaacaatgctaaacccaaacagttactgcatttttttagacag<br>ccctaactggtgttccctcttcacaggaactgactgatcatggctagcggatccggaagtggtccatcaagactcgaggaggagttgagacgaagactgacagagcctggatctggaagcg<br>gtccttctagacttgaggaggagctgaggagaagactgaccgagccaggatctgggtctggaccatctagactggaggaggagctgaggcgcaggctgaccgaacctggctctggctcc<br>ggaccaagcaggctggaagaggagctgagaagaaggctgactgaacctggaagtggttctggaccttctagactggaggaggagctgagaagacggctgaccgaaccaggcagtggc<br>tcaggcccatcaagactggaggaagagctgcgtcgtaggctgacagaaccaggctctggatcaggaccctctagacttgaagaggagctgagaagaagactgacagagcctggatctg<br>ggtctggcccttctagactggaagaggagctgagaagaagactgacagagcctggatctggatcaggtccttctagactggaggaggagctgagacggcggctgacagagccaggatct<br>ggaagcggacctagcaggctggaggaggagctgaggaggagactgaccgaaccaggatctggatctggacctagcaggctggaggaggagctcaggagacgtctgacagagccag<br>gaagcggaagcggaccatctagactggaggaggagctgagaaggcgtctgacagagcccggatccggctctggaccttctcgactcgaggaagagctgagaaggagattaacagagc<br>ccggcagcggatccggaccaagccgcctggaggaagagctgagacgaaggctgactgagcctggatctggttcaggaccctctaggctcgaggaggagctgagacgtaggctgactg<br>aacctggatctggatctggacctagccgcctggaggaggagctgcgcagaagactgaccgagcctggaagtggctccggaccatcaaggctggaggaggaactgagaagacgcctga<br>ctgagccaggatcaggctcaggacccagcaggctggaggaggagctgagaagaagactgaccgagcctggatccggatctggaccaagcagactggaagaggaactgagaagaag<br>gctgacagagcctggatcaggatcaggccccagcagactggaagaggagctgagaaggagactgacagagccaggatcaggatcaggcccttcaagactggaggaggagctgcgca<br>gaagactgacagagcccggatctggatccggcccaagtcgcctggaggaggagctgagaagaagactgacagagcctggaagcggttccggcccttctcggctggaggaggagctga<br>gaagaagactgaccgagcccggtagcggatctggcccaagcaggctggaagaggagctgcgtaggagactgactgagcccggatctggaagtggaccttctagactggaggaggagc<br>tgagaagaagactgactgaaccaggatcaggaagcggaccttcccgtctggaggaagagctgcgcaggcgactgactgagccagggtccggatcaggaccaagccgcctcgaggag<br>gagctcagaagaagactgacagagccaggatctggcagcggaccatctaggctggaggaggagctgaggcgcaggctgactgagcctggttccggaagcggaccatccagactgga<br>ggaggagctgagaaggagactgacagaacctggaagcggcagcggcccctctagattggaagaggagctgagaaggagactgactgagccaggctctggctccggaccaagccgcc<br>tggaggaggagctgaggagacggctgacggagcctggatcaggaagcggaccttctcgtctggaggaggagctgagaagacgactgacagagcccggatcaggcagtggcgacgg<br>cgctagcgtcgccgtggtccgtgctctcacggtgctgttgctcggtcaggtgttgctgggaggtgccgttggactcattcccgagatcgaccgacggaaatacagtgattcggggagacac<br>acaccggagcgaactgatacaaacttcctgaacgagtttgagctacgcttgctcaatatgttcggattgaagcgaaaacccacccccaagcaaatcggcagtggtccctcagtacatgctgga<br>cttgtattatatgcactctgaaaacgatgacccgaacattcggcgcccgaggagcactatgggaaaacatgtagaaagggcagccagcagagcaaacacgatacgaagttttcatcacgaa<br>ggtgaggcctgatttgatattacagcctgaaaatgtacacaattttccagtgtaaaagaatgtttgtttattcccggaacattccaaagttctataggaatgatgtgatgcaaccgtttggaaagca<br>gcgacattccacaagcggctctcactgttattaaagaagcaagaaatggcatcctaacggtcactttgtgcaaataattgatagtgtttgagtgcacttagagtagttttccagacaaggtgtgga<br>gtttatgcactagttattttccaagaatccgcagattgcagtaacaagccctacgaggaagtccaggatggtagttttgctgttctccatcaagttctgtcattgatgtttttgagccctcaatggga<br>gggcttgtttctgcaccaacttacagatattgtgagtgtgtgtgtgtgtttagctttgctcaactctttatcacatgcccgagtaactctagcctaacttgccagattgtgggtggccttgttgaatggt<br>cagatttaattcacaagaaactgtatccgcttttctttttgatacatataatgaaatcaaatccgagtgtgatcaggtggcatgtcaattaagtctcttttaggtgtgttatctgatttaaacaatggtga<br>aacatgccctctcttgctctggaagacttgagcctgtgaaagtcaccggcctcgttcatgcaagtcttccctctggagtctgtttagaggtcagtgtaaactgtagcgcacgatttagaaacattt<br>cactttagtttcagggatacacaggacaaaatcacgctaacttgctgtactgtggtaaagagagggggagggatgcatacgagagtcattacagggctttttaggagtaggctggctctgcaa<br>aatggcacagatttggatggatgacatccagtggttgagctgaaacactgcaaactacgatgcttcttgcgcactcgcacttcaacgcacaatctaagcatttgcgtgttccttgttgattccaaa<br>aggaaacttttctctatagttcctcctacgcctctggcatttctcattgatgtgatgaactctgtggttgagcacctgatgggaattttattacatcccattttggcacggcgttatgtaattgttttaaag<br>ttccagccaaaatggagcttttgaaatcacttgttttgatgtcagtcctcctgagagattctcagctgaggtgcgcagtcaagtagccgtcctacacgtacggcttgtacgatttgctgttttggcat<br>aaggtacaaacagaaattaggaatgtcaatacactcataactgtggctgactgcattcggcaagccccgacatgttttcagtcactcccccctcaagacggattgactgggagaaaaaggaaa<br>aggtgaagaaagacagacaaaaatgagggaaagaaagagctgggctttcccacatcaaataaagtgcgcttatcttctgtaggctctttgtgattatgcagcagataggttggattattgggaa<br>tggaatttccaaaacgactcttttcaaaagagattaagcgagctctttggtctgagagcgttttattttagggcattgagtcgtaggagtagctgtttcttcgctgattgcaacctttattcaataaag<br>acttggaaacaagagaatgttgtcactctacagccacacaaacagccttgtgcttacagctggcctttcattttaacacctggtcctggtaccttataatatccagcgcaggccagaccagcaaa<br>ttgctgagtggaaaatcccccccccaaaaaagacttgttagatgtttctaacaaactcttttcttctgtccttccagaggctttcgaggcactgtccagcctgaaaggaaaaacaacgcagcagttt<br>ttcttcaaccttacctccattcctggcgaggagctgatctccgctgcggagctgcgcattttcagggaccaagttctcggagatgccagtacgagtggcttccacagaattaacatttacgaggt<br>gttcaggccagctttggccccctccaaagagcctctaaccagacttctggacacccgtctggtgcaggactctcacacgcgctgggaaagcttcgacgtgggttcagctgtggcacgctgg<br>gcccgcgaatcccagcacaaccatgggctccttgtagaggtgctccatcctaaggagtcagaagtatccgaggaggctgagagcaaccggaggaagcacgtgagggtcagtcgttccct<br>tcacgcggatgaggactcgtgggcacaagcccgacctctgctggtaacctacagccatgacggtcaaggcacagccgtcttgcattcgaaccgagaaaagcggcaggctcgacgaggg<br>caaaagccgaggagaaagcaccaccagcgctcgaactgtaggcgacatgctctctatgtggacttcagtgatgtcggctggaacgagtggatcgtggcaccgccaggctatcatgctttct<br>actgccatggcgagtgtccgttccctctgccggaccatctaaactccaccaaccatgccattgtccagacgctggtgaactcggtcaactccaacattcccaaagcctgttgcatcccgacgg<br>agctcagccctatctcactgctgtacctggacgagtatgagaaggtcattcttaaaaactaccaggacatggtggtggagggctgcggttgccgatgacccggggaacaatctccccaatga<br>agacttttatttatacaaaagagcgagctattttggaggaagaaaagaaatatatatgaatatatttatgttgaatgaacaaaacaaaaaaaatgggaaaataaatattttaatgagagtgcttgctt<br>tttccagtggcttcgaagatgatttttttcttgccccattccaaaacaagtgcaatggcacatgaagtataatgctcagatttttatgatgtatttattgcataaccactttatttgtaaatgtgtatttatc<br>atgaaaaaaatatatgatcttcattcagtgtgcatttgggaataccttatttgataccaaaaaaaataaacaatggatgatgaa |
| ***bmp2b_ALFA_array***<br><br>bmp2b promoter (5'UTR) > ALFA_array - bmp2b (intron) – 30X MS2 - 3'UTR | cattgctcctcttcccagtttggtcgagggttaacataatatcaagtcgtgcatttcctattgacaatatagcatgtttacctgtggattgatgtataaaacaagttcgactgatttatgcaaagcaaa<br>cacactgttgtgttacagcaaccttctgacacatcagcaatcgatacacacagagcattgttatttaaatcagacttctccagaacagtgtctgctcaggatttaacttgatagtgtgaactgattgt<br>gatgcaagatgtgttggaaatatatatataaagtaatgtatacttttacactattttacagcacaattaaacagattttctattagcacaggggttttaaaactcccaagggccgtaaaaggggtgct<br>gtcttgtgtttttctatagagggaggctactgttgcctgtaggctatgtcttcaggaaaagggtagtttgaacattgaagtaaatattactatgaatttcattaggtgaaaatgtttcatgtgtaacattt<br>atacaagcaactgcactttaatattgcgggaaaatgtaaatgtgtttactattttaatactaaacatatgcaattataattgactctataaagactctataaattgttgtgagcatcacattttggacaaa<br>tcaaaagtatccggaaaatataatgcatacctgtacactttgagttcttttttttttttttgaacacaggagctttaaaactgagggccggggactcccaggagtccataaagtggtgccatcttttc<br>ttttcttctttttttcttctgcttgtctagccagtggcaataatctccctcaggcaatattttctgtaaagctgctaaagaaaaaagtggcatagaaataaatgttacatatatatatatacatacacagtg<br>ttcatcatatataagtacaccctttgaaaatctctcaattcatattttgtgcatatactttagataagtcagtactgaagccaaatctgaagctaatctaagaaaataacatgatagtctaagttagtac<br>actcaaatttatgtaagggaaaatattaagtaaaaaatgtaaaaagagcacaattcatatatgtatattgtttatatatgtatacatataggaagctttacaatattatatttgtgcatatacattagacta<br>gtcagtattaaagccaaatctggagctaatttaacaaaataacttatgattatagtttaaaatttgtacacgcaaatttataagttaagcaaatatatatatatatatatatatatatatacatacatac<br>atacatacatacctcaattcaaggtatttttatttatttatttatttttgttaactaactgtaaacagaattattagccccccttgtttatttttttcccccatttcttctgtttaacggaaagcagattttttaagc<br>agactttaaaatgtttctttaaaaataaaaaacttgttattttctagccaaaataaaacaaataagactttctcccttgctctgataaaaatcatttgggaaatataaaaaaaaagaacaaaatttcaaa<br>ggggcactaataattctgacatcaactttaaattttatttatttatcttttggtaactacgacaaaaagagatgtaatttagctttatagctaactctggcataacatgtcatgttgtagctacattgtccc<br>agaataagtaaataaatgaatattcggctttatacaagtctaaaatagttgtacataaaatattagatcattttaaaacgtttaaagacaacacattgcaataacaaatcaattaaatgaaacctaaa<br>ataacgttaacatttttccttcactataaattactatacatgattttaaacagaagagatatccttataaatactgaaaaaaatactcaaatacaaatgtagataatttaaattagtgcgcatttaaattta<br>ggatttgtttaaccatacttcagtctcaattgtattgcgtatacattacgttctcgttcaaattactaacatgtttacataggataatgcataaaatatgccccatgcaggggaaattcggtccatccgc |

| | |
|---|---|
| | gcgcgcagagtgtgggcatgttcaaacgcttgaatggagagagcgcggcatcattgtgacatcatcagacaacaaaaagcctcgcgctcgcgcagcgaagcgctccaatcaatggcaca<br>gacgcggcgcgtgctgcacgcagagatgagtctccaaacagccacggaaaacttctgctgaccacaagttttcgatttctttaaaacaaaaacaaaaaatgacaaatccaggattgtgcgat<br>ctcgcgctgtcacttttgggattgctgctgtctttgacctgagcgctcgcgcacttcattagagtttagtagagtctagtctgaagtgttgcacaagtatgaacaagaagaggcgacttgagctgc<br>gacgactctctgtcgtgggataaaaaaatcgcttgtggattaaaacacgaattcatgaggaacttagaagacgacgggaacgcagaccggccacagcgcttcctcctccggtaacgcattc<br>aattacatgcttgatattacgtattagcaaacagtgcgactgttttaaaaactaatgctgacatttctgtaactttgattgcacttcacatcactgcgtgtgcgtgcctgcatgtaatgtgtgaggtca<br>ggaggtttgcgcaaatgaaaagaaaacgcttctgcgcttccttatcatcagatctctttgtcgtctttgttgcccgagttgtttttgcgtaagatcgaccacctgtagcctcacaactctgcctctcc<br>cttagcgaaatgtcttttgctagacattcagtccttttgaagactttcctttcatgatctaaagacagacccccacccagacagcgcgctcttatctccctttaattgcagactccatgtctgctgtctt<br>catgtctcccttctttcgatttcttttgattcactctttcctgtttcatcccatttgtctgacatcctctttggagtctcccgctttcactgtagatcaatctcttattatagcctgaggcagggatattattctt<br>aatcttttttttagagcgcgagtgtgtgttcatcgcctcaaagacatctccaaaagatccagatatgtctgtggtttgaggcagccgcaggttgaacgcggccagggaacttttccatgcgtataa<br>gtggttgaagctctgggcttgaagtcaaacgtgtttttttaggacttaaaggcgcaggtattccataaaatcaaactattatgagttattggcggattgaacctatctttaggtgatctatgagccatt<br>gtgccactgagcaaggcacttattaacgacaccagattgctccaagaggggctgtagtttaaaaatgtactcttcaaaaataaagttattttaaataattaaactttacttacagaatatattttcttg<br>caatatttaaacattcttaaatcaagaacatcacagttttgtttttagaaataagaagtgaaattaagtgaggctttctttaaaacaagctaaattatttccagtggggtaagtaaataatcttatgtcaa<br>actaaaagcaagaattgtttactggcttgatttgaggaaaaacacttgatttcgacgttttatttctgaaaacaaagcatttcacttgtttagaaaatgcaattcgatagatagatatttgatttttagat<br>atttacactagaaaccagacaaaaaccattttaattttcaggaaatgcatgtttttatgcagcgctttacattatttttactgttacgttacaggtttagatcaatgtaaagttattatagttattggtggat<br>tggacctatcttttagatgatctatgagccattgtgccactaagcaaggcacttaattgctccagagattgctccagaagggctgtagatttaagaaatgtactcttaaaaataaacttattttatgaa<br>ttaatcttcacgaaaaatatatattcttactctcgagtctaaatatttaaacattgtaaaatcaagaagcattttctaaacaagtaaggtccaaattaagtgaggatttctttaaagcaagctcaattatc<br>ttccagtagggtaagtaaaataatcttatttcaaactaaaagcaagattattttactcaccaaactggttagatttgaggaaaaacacataatttttacattttatttctgaaagcaaaacattgcacttg<br>tctatgaataatatgatgtaaatataatttaataaattattaataatttaaattatgaaattgaataataatacaataataataaaatattaatatatatatatatttacacctgaaacgagacaaaaactctt<br>tttggaaaaccacactttatgcagtgctatcatgtattttaattaaaatgttatattacaagttcagatcgatgtgttgctttggattctgtctcccttaaagcagcagagcatgcttttcccaagaaacc<br>agaaaataaaaagttcactgtgggccctaaaaaggttcatctctatgatgcaatgctgtaaaacctttaaaatgatttatggtaagaacaatgctaaacccaaacagttactgcatttttttagacag<br>ccctaactggtgttccctcttcacaggaactgactgatcatggctagcggatccggaagtggtccatcaagactcgaggaggagttgagacgaagactgacagagcctggatctggaagcg<br>gtccttctagacttgaggaggagctgaggagaagactgaccgagccaggatctgggtctggaccatctagactggaggaggagctgaggcgcaggctgaccgaacctggctctggctcc<br>ggaccaagcaggctggaagaggagctgagaagaaggctgactgaacctggaagtggttctggaccttctagactggaggaggagctgagaagacggctgaccgaaccaggcagtggc<br>tcaggcccatcaagactggaggaagagctgcgtcgtaggctgacagaaccaggctctggatcaggaccctctagacttgaagaggagctgagaagaagactgacagagcctggatctg<br>ggtctggcccttctagactggaagaggagctgagaagaagactgacagagcctggatctggatcaggtccttctagactggaggaggagctgagacggcggctgacagagccaggatct<br>ggaagcggacctagcaggctggaggaggagctgaggaggagactgaccgaaccaggatctggatctggacctagcaggctggaggaggagctcaggagacgtctgacagagccag<br>gaagcggaagcggaccatctagactggaggaggagctgagaaggcgtctgacagagcccggatccggctctggaccttctcgactcgaggaagagctgagaaggagattaacagagc<br>ccggcagcggatccggaccaagccgcctggaggaagagctgagacgaaggctgactgagcctggatctggttcaggaccctctaggctcgaggaggagctgagacgtaggctgactg<br>aacctggatctggatctggacctagccgcctggaggaggagctgcgcagaagactgaccgagcctggaagtggctccggaccatcaaggctggaggaggaactgagaagacgcctga<br>ctgagccaggatcaggctcaggacccagcaggctggaggaggagctgagaagaagactgaccgagcctggatccggatctggaccaagcagactggaagaggaactgagaagaag<br>gctgacagagcctggatcaggatcaggccccagcagactggaagaggagctgagaaggagactgacagagccaggatcaggatcaggcccttcaagactggaggaggagctgcgca<br>gaagactgacagagcccggatctggatccggcccaagtcgcctggaggaggagctgagaagaagactgacagagcctggaagcggttccggcccttctcggctggaggaggagctga<br>gaagaagactgaccgagcccggtagcggatctggcccaagcaggctggaagaggagctgcgtaggagactgactgagcccggatctggaagtggaccttctagactggaggaggagc<br>tgagaagaagactgactgaaccaggatcaggaagcggaccttcccgtctggaggaagagctgcgcaggcgactgactgagccagggtccggatcaggaccaagccgcctcgaggag<br>gagctcagaagaagactgacagagccaggatctggcagcggaccatctaggctggaggaggagctgaggcgcaggctgactgagcctggttccggaagcggaccatccagactgga<br>ggaggagctgagaaggagactgacagaacctggaagcggcagcggcccctctagattggaagaggagctgagaaggagactgactgagccaggctctggctccggaccaagccgcc<br>tggaggaggagctgaggagacggctgacggagcctggatcaggaagcggaccttctcgtctggaggaggagctgagaagacgactgacagagcccggatcaggcagtggcgacgg<br>cgctagcgtcgccgtggtccgtgctctcacggtgctgttgctcggtcaggtgttgctgggaggtgccgttggactcattcccgagatcgaccgacggaaatacagtgattcggggagacac<br>acaccggagcgaactgatacaaacttcctgaacgagtttgagctacgcttgctcaatatgttcggattgaagcgaaaacccaccccaagcaaatcggcagtggtccctcagtacatgctgga<br>cttgtattatatgcactctgaaaacgatgacccgaacattcggcgcccgaggagcactatgggaaaacatgtagaaagggcagccagcagagcaaacacgatacgaagttttcatcacgaa<br>ggtgaggcctgatttgatattacagcctgaaaatgtacacaattttccagtgtaaaagaatgtttgtttattcccggaacattccaaagttctataggaatgatgtgatgcaaccgtttggaaagca<br>gcgacattccacaagcggctctcactgttattaaagaagcaagaaatggcatcctaacggtcactttgtgcaaataattgatagtgtttgagtgcacttagagtagttttccagacaaggtgtgga<br>gtttatgcactagttattttccaagaatccgcagattgcagtaacaagccctacgaggaagtccaggatggtagttttgctgttctccatcaagttctgtcattgatgtttttgagccctcaatggga<br>gggcttgtttctgcaccaacttacagatattgtgagtgtgtgtgtgtgtttagctttgctcaactctttatcacatgcccgagtaactctagcctaacttgccagattgtgggtggccttgttgaatggt<br>cagatttaattcacaagaaactgtatccgcttttctttttttgatacatataatgaaatcaaatccgagtgtgatcaggtggcatgtcaattaagtctcttttaggtgtgttatctgatttaaacaatggtga<br>aacatgccctctcttgctctggaagacttgagcctgtgaaagtcaccggcctcgttcatgcaagtcttccctctggagtctgtttagaggtcagtgtaaactgtagcgcacgatttagaaacattt<br>cactttagtttcagggatacacaggacaaaatcacgctaacttgctgtactgtggtaaagagagggggagggatgcatacgagagtcattacagggctttttaggagtaggctggctctgcaa<br>aatggcacagatttggatggatgacatccagtggttgagctgaaacactgcaaactacgatgcttcttgcgcactcgcacttcaacgcacaatctaagcatttgcgtgttccttgttgattccaaa<br>aggaaacttttctctatagttcctcctacgcctctggcatttctcattgatgtgatgaactctgtggttgagcacctgatgggaattttattacatcccattttggcacggcgttatgtaattgttttaaag<br>ttccagccaaaatggagcttttgaaatcacttgttttgatgtcagtcctcctgagagattctcagctgaggtgcgcagtcaagtagccgtcctacacgtacggcttgtacgatttgctgttttggcat<br>aaggtacaaacagaaattaggaatgtcaatacactcataactgtggctgactgcattcggcaagccccgacatgttttcagtcactccccctcaagacggattgactgggagaaaaaggaaa<br>aggtgaagaaagacagacaaaaatgagggaaagaaagagctgggctttcccacatcaaataaagtgcgcttatcttctgtaggctctttgtgattatgcagcagataggttggattattgggaa<br>tggaatttccaaaacgactcttttcaaaagagattaagcgagctctttggtctgagagcgttttattttagggcattgagtcgtaggagtagctgtttcttcgctgattgcaacctttattcaataaag<br>acttggaaacaagagaatgttgtcactctacagccacacaaacagccttgtgcttacagctggcctttcattttaacacctggtcctggtaccttataatatccagcgcaggccagaccagcaaa<br>ttgctgagtggaaaatcccccccccaaaaaagacttgttagatgtttctaacaaactcttttcttctgtccttccagaggctttcgaggcactgtccagcctgaaaggaaaaacaacgcagcagttt<br>ttcttcaaccttacctccattcctggcgaggagctgatctccgctgcggagctgcgcattttcagggaccaagttctcggagatgccagtacgagtggcttccacagaattaacatttacgaggt<br>gttcaggccagctttggccccctccaaagagcctctaaccagacttctggacacccgtctggtgcaggactctcacacgcgctgggaaagcttcgacgtgggttcagctgtggcacgctgg<br>gcccgcgaatcccagcacaaccatgggctccttgtagaggtgctccatcctaaggagtcagaagtatccgaggaggctgagagcaaccggaggaagcacgtgagggtcagtcgttccct<br>tcacgcggatgaggactcgtgggcacaagcccgacctctgctggtaacctacagccatgacggtcaaggcacagccgtcttgcattcgaaccgagaaaagcggcaggctcgacgaggg<br>caaaagccgaggagaaagcaccaccagcgctcgaactgtaggcgacatgctctctatgtggacttcagtgatgtcggctggaacgagtggatcgtggcaccgccaggctatcatgctttct<br>actgccatggcgagtgtccgttccctctgccggaccatctaaactccaccaaccatgccattgtccagacgctggtgaactcggtcaactccaacattcccaaagcctgttgcatcccgacgg<br>agctcagccctatctcactgctgtacctggacgagtatgagaaggtcattcttaaaaactaccaggacatggtggtggagggctgcggttgccgatgattgtaggcatgagtacagcctagaa<br>cggatacgattacgatccggcacatacacgaatacggtatgatctgcgtaccactaggacgcaacggtgccagcagtagcggcacagaatcatatcactagaatgattacccttcagaattat<br>cgaaggtgcggacgacaaatatgcgtcctctagtgcatcattagagcactctcctaagatgaatacacttagctatcgttactactaagaacgacgtaaaggaccaataggcctttctatttcta<br>gcactagcagaaagccatggtaggactaccaccatttgtaaacaccattagggtttatgcgctcgagcaatagccgagctatcgctgacgagtacgcagcgggatgactagaagtatcagtc<br>aagtacatgaggattacccatgttatcagagagaaatatcctctgaatgtttgactactaagcaaacgcttccatagtattaacatggaccggaagcatcagtagagcttctctgcctcaggagt<br>accgaggccctgacccagtagtaacgggtcgcgaccgcacaattatggcggtacccttgcagcattagcgcaagcgacttggagtaataacccaaggcgccgtcatcaatagagacggct<br>aaggctatgagtacaagcctcccggggaacaatctccccaatgaagacttttatttatacaaaagagcgagctattttggaggaagaaaagaaatatatatgaatatatttatgttgaatgaaca<br>aaacaaaaaaaatgggaaaataaatattttaatgagagtgcttgctttttccagtggcttcgaagatgatttttttcttgccccattccaaaacaagtgcaatggcacatgaagtataatgctcagat<br>ttttatgatgtatttattgcataaccactttatttgtaaatgtgtatttatcatgaaaaaatatatgatcttcattcagtgtgcatttgggaataccttatttgataccaaaaaaataaacaatggatgatga<br>a |
| ***bmp2b* 5'UTR for *in vitro* transcription** | taatacgactcactatagggcgtgctgcacgcagagatgagtctccaaacagccacggaaaacttctgctgaccacaagttttcgatttctttaaaacaaaaacaaaaaatgacaaatccag<br>gattgtgcgatctcgcgctgtcacttttgggattgctgctgtctttgacctgagcgctcgcgcacttcattagagtttagtagagtctagtctgaagtgttgcacaagtatgaacaagaagaggcg<br>acttgagctgcgacgactctctgtcgtgggataaaaaaatcgcttgtggattaaaacacgaattcatgaggaacttagaagacgacgggaacgcagaccggccacagcgcttcctcctccg<br>gaactgactgatcatggctagcggcagcggctcagggccaagccgcctcgaggaagagcttcgtagaagactgacagaacctgggtcgggttccggtccatcacgcttggaagaggag |

| | |
|---|---|
| T7 promoter > *bmp2b* (5'UTR) - ALFA_array - bmp2b - 3'UTR | ctgcgaagacggctgaccgaaccgggttctggtagcggacccagcaggctggaagaggaactgcgtcgacgcctcaccgagccaggctcgggaagcggtcccagtcgactggagga<br>ggaacttcgtcgccgacttacagaaccaggatcaggttccggcccatctcgcctggaggaggagctgcggcggcgtctcaccgaaccgggttccgggtccggtccgtcgcgcctggag<br>gaagaattgcggcgccgactaacggaaccgggctccggcagcggaccaagtcgcttggaggaggaacttcgccgtagactcactgaaccaggatctggatcaggaccatcgcgcttag<br>aagaagaactgcgccgcaggctgaccgagcccggctcgggatcgggcccatcgcgtctggaggaggagctgcggcgccgcttaacagagccaggtagtggcagcggcccctcccgc<br>ctggaagaggagctgcgcagaaggttgacggaacccggctctggctcaggcccatcccgactcgaagaggaattgcggcgtcgactgacggaaccgggctccggatccggacccagt<br>aggctagaagaagagctgcggcggcgtctcacagagccaggcagcggcagcggtccctcgcgcctggaagaggaactgcgacgtcgcttaaccgaaccgggatccggatctggacct<br>tcgcgcttggaggaggaactgcgtcgccgcctgacggaaccgggcagcggctccggaccctcaaggttggaggaggagctccgccggcgactgaccgagcccggcagtggatcagg<br>accatcgcgcctggaagaggagctgaggcggcggctgaccgaacctggtagcggcagcggcccctcccgtctagaggaagagctaaggcgtcgtcttaccgagccgggcagtggctc<br>gggtcccagcagactcgaggaggagttacgacgccgtctaactgagcctggttcagggagtggaccgtccaggctcgaggaggagctcaggcggaggctgacagagccgggaagtg<br>gctccggtccttcgcgcctggaggaagagctgcgccggcgccttacggagccaggcagcggcagcggacctagtcgcctggaagaagagctgagacgacggctgactgagccgggc<br>agcggctccggtccaagccgtttggaagaggagttaaggcgtcgcctcacggagcccggatccggctccggcccctctcgtttggaggaggagctgaggcgacggctcactgaacccg<br>ggtccggcagcggtccgtcccggctagaagaggagctgcgacgccgccttactgaaccagggagtggctctggaccctcgcgattggaggaggagctgcgccggcgcttgaccgagc<br>caggttcgggatcgggaccgagtcgcctggaggaggagctccggcgacgattgacggagccggggagcgggtccggtccttcacgcctggaggaggagttgcgcagacgcttgactg<br>aacccggatccggcagtgggccgagccgattggaggaggagcttcgacgccgactaaccgaacccggaagcggttcgggccccagccgcctggaggaagagctccggcgccgact<br>gacagagcccggatctggtagtggcccgtcgcgcttggaggaggagttgcgtcgtaggctgaccgaacccggctccggcagcggtccgtctagacttgaggaagaactgaggaggaga<br>ttgaccgagcctggctcgggctccggaccgtcacgattggaggaggagctaagacgcagacttacggagcccggatcgggcagcggtgctagcgtcgccgtggtccgtgctctcacggt<br>gctgttgctcggtcaggtgttgctgggaggtgccgttggactcattcccgagatcgaccgacggaaatacagtgattcggggagacacacaccggagcgaactgatacaaacttcctgaac<br>gagtttgagctacgcttgctcaatatgttcggattgaagcgaaaacccaccccaagcaaatcggcagtggtccctcagtacatgctggacttgtattatatgcactctgaaaacgatgacccga<br>acattcggcgcccgaggagcactatgggaaaacatgtagaaagggcagccagcagagcaaacacgatacgaagttttcatcacgaagaggctttcgaggcactgtccagcctgaaagga<br>aaaacaacgcagcagtttttcttcaaccttacctccattcctggcgaggagctgatctccgctgcggagctgcgcattttcagggaccaagttctcggagatgccagtacgagtggcttccaca<br>gaattaacatttacgaggtgttcaggccagctttggccccctccaaagagcctctaaccagacttctggacacccgtctggtgcaggactctcacacgcgctgggaaagcttcgacgtgggtt<br>cagctgtggcacgctgggcccgcgaatcccagcacaaccatgggctccttgtagaggtgctccatcctaaggagtcagaagtatccgaggaggctgagagcaaccggaggaagcacgt<br>gagggtcagtcgttcccttcacgcggatgaggactcgtgggcacaagcccgacctctgctggtaacctacagccatgacggtcaaggcacagccgtcttgcattcgaaccgagaaaagcg<br>gcaggctcgacgagggcaaaagccgaggagaaagcaccaccagcgctcgaactgtaggcgacatgctctctatgtggacttcagtgatgtcggctggaacgagtggatcgtggcaccg<br>ccaggctatcatgctttctactgccatggcgagtgtccgttccctctgccggaccatctaaactccaccaaccatgccattgtccagacgctggtgaactcggtcaactccaacattcccaaag<br>cctgttgcatcccgacggagctcagccctatctcactgctgtacctggacgagtacgagaaggtcattcttaaaaactaccaggacatggtggtggagggctgcggttgccgatgacccgg<br>ggaacaatctccccaatgaagacttttatttatacaaaagagcgagctattttggaggaagaaaagaaatatatatgaatatatttatgttgaatgaacaaaacaaaaaaaatgggaaaataaata<br>ttttaatgagagtgcttgctttttccagtggcttcgaagatgattttttttcttgccccattccaaaacaagtgcaatggcacatgaagtataatgctcagatttttatgatgtatttattgcataaccactt<br>tatttgtaaatgtgtatttatcatgaaaaaaatatatgatcttcattcagtgtgcatttgggaataccttatttgataccaaaaaaaataaacaatggatgatgaa |
| **ZIKV 5'UTR for *in vitro* transcription**<br><br>T7 promoter > ZIKV 5'UTR- ALFA_array - bmp2b - 3'UTR | taatacgactcactatagggagttgttgatctgtgtgaatcagactgcgacagttcgagtttgaagcgaaagctagcaacagtatcaacaggttttattttggatttggaaacgagagtttctggtc<br>atggctagcggcagcggctcagggccaagccgcctcgaggaagagcttcgtagaagactgacagaacctgggtcgggttccggtccatcacgcttggaagaggagctgcgaagacgg<br>ctgaccgaaccgggttctggtagcggacccagcaggctggaagaggaactgcgtcgacgcctcaccgagccaggctcgggaagcggtcccagtcgactggaggaggaacttcgtcgc<br>cgacttacagaaccaggatcaggttccggcccatctcgcctggaggaggagctgcggcggcgtctcaccgaaccgggttccgggtccggtccgtcgcgcctggaggaagaattgcggc<br>gccgactaacggaaccgggctccggcagcggaccaagtcgcttggaggaggaacttcgccgtagactcactgaaccaggatctggatcaggaccatcgcgcttagaagaagaactgcg<br>ccgcaggctgaccgagcccggctcgggatcgggcccatcgcgtctggaggaggagctgcggcgccgcttaacagagccaggtagtggcagcggcccctcccgcctggaagaggag<br>ctgcgcagaaggttgacggaacccggctctggctcaggcccatcccgactcgaagaggaattgcggcgtcgactgacggaaccgggctccggatccggacccagtaggctagaagaa<br>gagctgcggcggcgtctcacagagccaggcagcggcagcggtccctcgcgcctggaagaggaactgcgacgtcgcttaaccgaaccgggatccggatctggaccttcgcgcttggag<br>gaggaactgcgtcgccgcctgacggaaccgggcagcggctccggaccctcaaggttggaggaggagctccgccggcgactgaccgagcccggcagtggatcaggaccatcgcgcct<br>ggaagaggagctgaggcggcggctgaccgaacctggtagcggcagcggcccctcccgtctagaggaagagctaaggcgtcgtcttaccgagccgggcagtggctcgggtcccagca<br>gactcgaggaggagttacgacgccgtctaactgagcctggttcagggagtggaccgtccaggctcgaggaggagctcaggcggaggctgacagagccgggaagtggctccggtccttc<br>gcgcctggaggaagagctgcgccggcgccttacggagccaggcagcggcagcggacctagtcgcctggaagaagagctgagacgacggctgactgagccgggcagcggctccggt<br>ccaagccgtttggaagaggagttaaggcgtcgcctcacggagcccggatccggctccggcccctctcgtttggaggaggagctgaggcgacggctcactgaacccgggtccggcagcg<br>gtccgtcccggctagaagaggagctgcgacgccgccttactgaaccagggagtggctctggaccctcgcgattggaggaggagctgcgccggcgcttgaccgagccaggttcgggatc<br>gggaccgagtcgcctggaggaggagctccggcgacgattgacggagccggggagcgggtccggtccttcacgcctggaggaggagttgcgcagacgcttgactgaacccggatccg<br>gcagtgggccgagccgattggaggaggagcttcgacgccgactaaccgaacccggaagcggttcgggccccagccgcctggaggaagagctccggcgccgactgacagagcccgg<br>atctggtagtggcccgtcgcgcttggaggaggagttgcgtcgtaggctgaccgaacccggctccggcagcggtccgtctagacttgaggaagaactgaggaggagattgaccgagcctg<br>gctcgggctccggaccgtcacgattggaggaggagctaagacgcagacttacggagcccggatcgggcagcggtgctagcgtcgccgtggtccgtgctctcacggtgctgttgctcggt<br>caggtgttgctgggaggtgccgttggactcattcccgagatcgaccgacggaaatacagtgattcggggagacacacaccggagcgaactgatacaaacttcctgaacgagtttgagctac<br>gcttgctcaatatgttcggattgaagcgaaaacccaccccaagcaaatcggcagtggtccctcagtacatgctggacttgtattatatgcactctgaaaacgatgacccgaacattcggcgcc<br>cgaggagcactatgggaaaacatgtagaaagggcagccagcagagcaaacacgatacgaagttttcatcacgaagaggctttcgaggcactgtccagcctgaaaggaaaaacaacgca<br>gcagtttttcttcaaccttacctccattcctggcgaggagctgatctccgctgcggagctgcgcattttcagggaccaagttctcggagatgccagtacgagtggcttccacagaattaacattta<br>cgaggtgttcaggccagctttggccccctccaaagagcctctaaccagacttctggacacccgtctggtgcaggactctcacacgcgctgggaaagcttcgacgtgggttcagctgtggca<br>cgctgggcccgcgaatcccagcacaaccatgggctccttgtagaggtgctccatcctaaggagtcagaagtatccgaggaggctgagagcaaccggaggaagcacgtgagggtcagtc<br>gttcccttcacgcggatgaggactcgtgggcacaagcccgacctctgctggtaacctacagccatgacggtcaaggcacagccgtcttgcattcgaaccgagaaaagcggcaggctcga<br>cgagggcaaaagccgaggagaaagcaccaccagcgctcgaactgtaggcgacatgctctctatgtggacttcagtgatgtcggctggaacgagtggatcgtggcaccgccaggctatca<br>tgctttctactgccatggcgagtgtccgttccctctgccggaccatctaaactccaccaaccatgccattgtccagacgctggtgaactcggtcaactccaacattcccaaagcctgttgcatcc<br>cgacggagctcagccctatctcactgctgtacctggacgagtacgagaaggtcattcttaaaaactaccaggacatggtggtggagggctgcggttgccgatgacccggggaacaatctcc<br>ccaatgaagacttttatttatacaaaagagcgagctattttggaggaagaaaagaaatatatatgaatatatttatgttgaatgaacaaaacaaaaaaaatgggaaaataaatattttaatgagagt<br>gcttgctttttccagtggcttcgaagatgattttttttcttgccccattccaaaacaagtgcaatggcacatgaagtataatgctcagatttttatgatgtatttattgcataaccactttatttgtaaatgt<br>gtatttatcatgaaaaaaatatatgatcttcattcagtgtgcatttgggaataccttatttgataccaaaaaaaataaacaatggatgatgaa |
| **ZIKV 3'UTR for *in vitro* transcription**<br><br>T7 promoter > *bmp2b* (5'UTR) - ALFA_array - bmp2b - ZIKV 3'UTR | taatacgactcactatagggcgtgctgcacgcagagatgagtctccaaacagccacggaaaacttctgctgaccacaagttttcgatttctttaaaacaaaaacaaaaaatgacaaatccag<br>gattgtgcgatctcgcgctgtcacttttgggattgctgctgtctttgacctgagcgctcgcgcacttcattagagtttagtagagtctagtctgaagtgttgcacaagtatgaacaagaagaggcg<br>acttgagctgcgacgactctctgtcgtgggataaaaaaatcgcttgtggattaaaacacgaattcatgaggaacttagaagacgacgggaacgcagaccggccacagcgcttcctcctccg<br>gaactgactgatcatggctagcggcagcggctcagggccaagccgcctcgaggaagagcttcgtagaagactgacagaacctgggtcgggttccggtccatcacgcttggaagaggag<br>ctgcgaagacggctgaccgaaccgggttctggtagcggacccagcaggctggaagaggaactgcgtcgacgcctcaccgagccaggctcgggaagcggtcccagtcgactggagga<br>ggaacttcgtcgccgacttacagaaccaggatcaggttccggcccatctcgcctggaggaggagctgcggcggcgtctcaccgaaccgggttccgggtccggtccgtcgcgcctggag<br>gaagaattgcggcgccgactaacggaaccgggctccggcagcggaccaagtcgcttggaggaggaacttcgccgtagactcactgaaccaggatctggatcaggaccatcgcgcttag<br>aagaagaactgcgccgcaggctgaccgagcccggctcgggatcgggcccatcgcgtctggaggaggagctgcggcgccgcttaacagagccaggtagtggcagcggcccctcccgc<br>ctggaagaggagctgcgcagaaggttgacggaacccggctctggctcaggcccatcccgactcgaagaggaattgcggcgtcgactgacggaaccgggctccggatccggacccagt<br>aggctagaagaagagctgcggcggcgtctcacagagccaggcagcggcagcggtccctcgcgcctggaagaggaactgcgacgtcgcttaaccgaaccgggatccggatctggacct<br>tcgcgcttggaggaggaactgcgtcgccgcctgacggaaccgggcagcggctccggaccctcaaggttggaggaggagctccgccggcgactgaccgagcccggcagtggatcagg<br>accatcgcgcctggaagaggagctgaggcggcggctgaccgaacctggtagcggcagcggcccctcccgtctagaggaagagctaaggcgtcgtcttaccgagccgggcagtggctc<br>gggtcccagcagactcgaggaggagttacgacgccgtctaactgagcctggttcagggagtggaccgtccaggctcgaggaggagctcaggcggaggctgacagagccgggaagtg<br>gctccggtccttcgcgcctggaggaagagctgcgccggcgccttacggagccaggcagcggcagcggacctagtcgcctggaagaagagctgagacgacggctgactgagccgggc<br>agcggctccggtccaagccgtttggaagaggagttaaggcgtcgcctcacggagcccggatccggctccggcccctctcgtttggaggaggagctgaggcgacggctcactgaacccg |

| | |
|---|---|
| | ggtccggcagcggtccgtcccggctagaagaggagctgcgacgccgccttactgaaccagggagtggctctggaccctcgcgattggaggaggagctgcgccggcgcttgaccgagc<br>caggttcgggatcgggaccgagtcgcctggaggaggagctccggcgacgattgacggagccggggagcgggtccggtccttcacgcctggaggaggagttgcgcagacgcttgactg<br>aacccggatccggcagtgggccgagccgattggaggaggagcttcgacgccgactaaccgaacccggaagcggttcgggccccagccgcctggaggaagagctccggcgccgact<br>gacagagcccggatctggtagtggcccgtcgcgcttggaggaggagttgcgtcgtaggctgaccgaacccggctccggcagcggtccgtctagacttgaggaagaactgaggaggaga<br>ttgaccgagcctggctcgggctccggaccgtcacgattggaggaggagctaagacgcagacttacggagcccggatcgggcagcggtgctagcgtcgccgtggtccgtgctctcacggt<br>gctgttgctcggtcaggtgttgctgggaggtgccgttggactcattcccgagatcgaccgacggaaatacagtgattcggggagacacacaccggagcgaactgatacaaacttcctgaac<br>gagtttgagctacgcttgctcaatatgttcggattgaagcgaaaacccaccccaagcaaatcggcagtggtccctcagtacatgctggacttgtattatatgcactctgaaaacgatgacccga<br>acattcggcgcccgaggagcactatgggaaaacatgtagaaagggcagccagcagagcaaacacgatacgaagttttcatcacgaagaggctttcgaggcactgtccagcctgaaagga<br>aaaacaacgcagcagtttttcttcaaccttacctccattcctggcgaggagctgatctccgctgcggagctgcgcattttcagggaccaagttctcggagatgccagtacgagtggcttccaca<br>gaattaacatttacgaggtgttcaggccagctttggccccctccaaagagcctctaaccagacttctggacacccgtctggtgcaggactctcacacgcgctgggaaagcttcgacgtgggtt<br>cagctgtggcacgctgggcccgcgaatcccagcacaaccatgggctccttgtagaggtgctccatcctaaggagtcagaagtatccgaggaggctgagagcaaccggaggaagcacgt<br>gagggtcagtcgttcccttcacgcggatgaggactcgtgggcacaagcccgacctctgctggtaacctacagccatgacggtcaaggcacagccgtcttgcattcgaaccgagaaaagcg<br>gcaggctcgacgagggcaaaagccgaggagaaagcaccaccagcgctcgaactgtaggcgacatgctctctatgtggacttcagtgatgtcggctggaacgagtggatcgtggcaccg<br>ccaggctatcatgctttctactgccatggcgagtgtccgttccctctgccggaccatctaaactccaccaaccatgccattgtccagacgctggtgaactcggtcaactccaacattcccaaag<br>cctgttgcatcccgacggagctcagccctatctcactgctgtacctggacgagtacgagaaggtcattcttaaaaactaccaggacatggtggtggagggctgcggttgccgatgacccgg<br>gggcaccaatcttaatgttgtcaggcctgctagtcagccacagcttggggaaagctgtgcagcctgtgaccccccccaggagaagctgggaaaccaagcctatagtcaggccgagaacgc<br>catggcacggaagaagccatgctgcctgtgagcccctcagaggacactgagtcaaaaaaccccacgcgcttggaggcgcaggatgggaaaagaaggtggcgaccttccccacccttca<br>atctggggcctgaactggagatcagctgtggatctccagaagagggactagtggttagaggagacccccccggaaaacgcaaaacagcatattgacgctgggaaagaccagagactccat<br>gagtttccaccacgctggccgccaggcacagatcgccgaatagcggcggccggtgtggggaaatccatgggtct |
| ***actb2* 5’UTR for *in vitro* transcription**<br><br>T7 promoter > actb2 5’UTR - ALFA_array - bmp2b - 3’UTR | taatacgactcactatagggaagagcgcgcccagcttttcagcctcactttgagctcctccacacgcagctagtgcggaatatcatctgcttgtaacccattctcttaagtcgacaaccccccaa<br>acccaagttcagccatggctagcggcagcggctcagggccaagccgcctcgaggaagagcttcgtagaagactgacagaacctgggtcgggttccggtccatcacgcttggaagagga<br>gctgcgaagacggctgaccgaaccgggttctggtagcggacccagcaggctggaagaggaactgcgtcgacgcctcaccgagccaggctcgggaagcggtcccagtcgactggagg<br>aggaacttcgtcgccgacttacagaaccaggatcaggttccggcccatctcgcctggaggaggagctgcggcggcgtctcaccgaaccgggttccgggtccggtccgtcgcgcctggag<br>gaagaattgcggcgccgactaacggaaccgggctccggcagcggaccaagtcgcttggaggaggaacttcgccgtagactcactgaaccaggatctggatcaggaccatcgcgcttag<br>aagaagaactgcgccgcaggctgaccgagcccggctcgggatcgggcccatcgcgtctggaggaggagctgcggcgccgcttaacagagccaggtagtggcagcggcccctcccgc<br>ctggaagaggagctgcgcagaaggttgacggaacccggctctggctcaggcccatcccgactcgaagaggaattgcggcgtcgactgacggaaccgggctccggatccggacccagt<br>aggctagaagaagagctgcggcggcgtctcacagagccaggcagcggcagcggtccctcgcgcctggaagaggaactgcgacgtcgcttaaccgaaccgggatccggatctggacct<br>tcgcgcttggaggaggaactgcgtcgccgcctgacggaaccgggcagcggctccggaccctcaaggttggaggaggagctccgccggcgactgaccgagcccggcagtggatcagg<br>accatcgcgcctggaagaggagctgaggcggcggctgaccgaacctggtagcggcagcggcccctcccgtctagaggaagagctaaggcgtcgtcttaccgagccgggcagtggctc<br>gggtcccagcagactcgaggaggagttacgacgccgtctaactgagcctggttcagggagtggaccgtccaggctcgaggaggagctcaggcggaggctgacagagccgggaagtg<br>gctccggtccttcgcgcctggaggaagagctgcgccggcgccttacggagccaggcagcggcagcggacctagtcgcctggaagaagagctgagacgacggctgactgagccgggc<br>agcggctccggtccaagccgtttggaagaggagttaaggcgtcgcctcacggagcccggatccggctccggcccctctcgtttggaggaggagctgaggcgacggctcactgaacccg<br>ggtccggcagcggtccgtcccggctagaagaggagctgcgacgccgccttactgaaccagggagtggctctggaccctcgcgattggaggaggagctgcgccggcgcttgaccgagc<br>caggttcgggatcgggaccgagtcgcctggaggaggagctccggcgacgattgacggagccggggagcgggtccggtccttcacgcctggaggaggagttgcgcagacgcttgactg<br>aacccggatccggcagtgggccgagccgattggaggaggagcttcgacgccgactaaccgaacccggaagcggttcgggccccagccgcctggaggaagagctccggcgccgact<br>gacagagcccggatctggtagtggcccgtcgcgcttggaggaggagttgcgtcgtaggctgaccgaacccggctccggcagcggtccgtctagacttgaggaagaactgaggaggaga<br>ttgaccgagcctggctcgggctccggaccgtcacgattggaggaggagctaagacgcagacttacggagcccggatcgggcagcggtgctagcgtcgccgtggtccgtgctctcacggt<br>gctgttgctcggtcaggtgttgctgggaggtgccgttggactcattcccgagatcgaccgacggaaatacagtgattcggggagacacacaccggagcgaactgatacaaacttcctgaac<br>gagtttgagctacgcttgctcaatatgttcggattgaagcgaaaacccaccccaagcaaatcggcagtggtccctcagtacatgctggacttgtattatatgcactctgaaaacgatgacccga<br>acattcggcgcccgaggagcactatgggaaaacatgtagaaagggcagccagcagagcaaacacgatacgaagttttcatcacgaagaggctttcgaggcactgtccagcctgaaagga<br>aaaacaacgcagcagtttttcttcaaccttacctccattcctggcgaggagctgatctccgctgcggagctgcgcattttcagggaccaagttctcggagatgccagtacgagtggcttccaca<br>gaattaacatttacgaggtgttcaggccagctttggccccctccaaagagcctctaaccagacttctggacacccgtctggtgcaggactctcacacgcgctgggaaagcttcgacgtgggtt<br>cagctgtggcacgctgggcccgcgaatcccagcacaaccatgggctccttgtagaggtgctccatcctaaggagtcagaagtatccgaggaggctgagagcaaccggaggaagcacgt<br>gagggtcagtcgttcccttcacgcggatgaggactcgtgggcacaagcccgacctctgctggtaacctacagccatgacggtcaaggcacagccgtcttgcattcgaaccgagaaaagcg<br>gcaggctcgacgagggcaaaagccgaggagaaagcaccaccagcgctcgaactgtaggcgacatgctctctatgtggacttcagtgatgtcggctggaacgagtggatcgtggcaccg<br>ccaggctatcatgctttctactgccatggcgagtgtccgttccctctgccggaccatctaaactccaccaaccatgccattgtccagacgctggtgaactcggtcaactccaacattcccaaag<br>cctgttgcatcccgacggagctcagccctatctcactgctgtacctggacgagtatgagaaggtcattcttaaaaactaccaggacatggtggtggagggctgcggttgccgatgacccggg<br>gaacaatctccccaatgaagacttttatttatacaaaagagcgagctattttggaggaagaaaagaaatatatatgaatatatttatgttgaatgaacaaaacaaaaaaaatgggaaaataaatat<br>tttaatgagagtgcttgctttttccagtggcttcgaagatgattttttcttgccccattccaaaacaagtgcaatggcacatgaagtataatgctcagatttttatgatgtatttattgcataaccacttt<br>atttgtaaatgtgtatttatcatgaaaaaatatatgatcttcattcagtgtgcatttgggaataccttatttgataccaaaaaaataaacaatggatgatgaa |

**Table S1.**

Sequence of the plasmids used in this study.

**Movie S1.**

Maximum intensity projected light sheet movies of *Tg(ubi:NB-ALFA-msGFP2_HIGH)* embryos with maternal and zygotic expression (left) and zygotic expression only (right). Timing corresponds to after fertilization. Related to Fig. 1 and fig. S1.

**Movie S2.**

Maximum intensity projected LLS movie of a ~4.3 hpf *Tg(ubi:NB-ALFA-msGFP2_HIGH)* embryo injected with the *ALFA_array_bmp2b* plasmid. Related to Fig. 3C and fig. S3B.

**Movie S3.**

LLS movies of 1 hpf *Tg(ubi:NB-ALFA-msGFP2_HIGH)* embryos injected with *ALFA_array_bmp2b* mRNA without (top) or with puromycin (middle) or with *bmp2b* mRNA (no ALFA_array, bottom). Related to Fig. 2D.

**Movie S4.**

LLS movie of a ~4.3 hpf *Tg(ubi:NB-ALFA-mScarlet3); Tg(actb2:2xMCP-NLS-EGFP)* embryo injected with *ALFA_array_bmp2b_30xMS2* DNA. MCP-EGFP is shown in magenta and corresponds to mRNA molecules and NB-ALFA-mScarlet3 (nascent translation) is shown in cyan. Related to Fig. 2E.

**Movie S5.**

LLS movie of a ~4.3 hpf *Tg(ubi:NB-ALFA-mScarlet3); Tg(actb2:2xMCP-NLS-EGFP)* embryo injected with *ALFA_array_bmp2b_30xMS2* DNA. The movie shows a single molecule of mRNA (magenta) in translation (cyan). Related to Fig. 2E.

**Movie S6.**

Maximum intensity projected LLS movie of a *Tg(ubi:NB-ALFA-msGFP2_HIGH); Tg(bmp2b:ALFA_array_bmp2b)* embryo. Related to Fig. 3E.
